# Characterizing nascent readthrough transcripts in human cells

**DOI:** 10.64898/2026.11.01.763429

**Authors:** Ariel McShane, Ishwarya Venkata Narayanan, Michelle T. Paulsen, Hailey Blinkiewicz, Mario Ashaka, Brian Magnuson, Karan Bedi, Nina T. Yang, Jayendra Prasad, Jeffrey N. Dudley, Sethuramasundaram Pitchiaya, Emily Seiden, Patrick J. Grohar, Erez Liberman Aiden, Thomas E. Wilson, Mats Ljungman

## Abstract

Termination of transcription by RNA polymerase II (RNAPII) is critical to appropriately define transcriptome and proteome identity; yet mechanisms of termination regulation are incompletely understood. Current models suggest critical roles for the exonuclease XRN2, which is posited to “torpedo” out RNAPII after degrading uncapped and RNAPII-tethered nascent transcripts and posit an “allosteric model” wherein elongating RNAPII loses its processivity downstream of the transcription end site (TES) via unknown mechanisms. Here, we used nascent RNA Bru-seq to interrogate transcription termination across 16 human cell lines of distinct lineages, as part of the ENCODE 4 project, to identify key regulatory modes. We find that >75% of protein-coding genes engage in readthrough transcription under homeostatic conditions. Exposing human cells to oxidative stress or hyperosmotic conditions or splicing inhibitors cause an increase in readthrough transcripts globally. Using an auxin-inducible degradation approach we find that depletion of BRD4, MED14 or XRN2 increases the extent and pervasiveness of readthrough transcripts while depletion of SUPT16H results in shorter readthrough transcripts. Our study provides a comprehensive view of homeostatic readthrough transcripts in human cells and delineates key components regulating transcription termination.

## INTRODUCTION

Transcription by RNA polymerase II (RNAPII) occurs in three distinct phases: initiation, elongation, and termination^1^. Transcription initiation is highly regulated by the binding of pioneer transcription factors, remodeling of chromatin to form nucleosome-depleted regions, and binding of RNAPII and general transcription factors to form a preinitiation complex at promoter elements^2^. Following initiation, RNAPII promoter-proximal pausing and productive elongation into the gene body occur regulated by elongation factors, histone modifications, and RNAPII carboxy-terminal domain (CTD) phosphorylation events^3,4^. When reaching the ends of genes, 3’-end processing occurs upon specific modifications of RNAPII slowing its elongation, recognition of a polyadenylation site, recruitment of the cleavage and polyadenylation complex followed by polyadenylation of the 3’-end of the nascent transcript by poly(A) polymerase^2,5–9^. Following 3’-end cleavage of the nascent mRNA, the RNAPII continues elongation and synthesizes RNA until it is “chased down” by the 5’-3’ exonuclease XRN2 in what has been called the “torpedo model“^10–12^. The digestion of the cleaved RNA by the XRN2 exonuclease is very rapid and these readthrough transcripts accumulate only after XRN2 is ellminated^13–15^.

Thus, transcripts that are detected past the TES in the presence of XRN2 are predominantly transcripts that did not undergo 3’-end cleavage^13–15^.

It appears that termination of transcription past the TES is a stochastic event rather than each gene having a preferred termination site. Genomic features that promote transcription termination include R-loops, which are enriched over G-rich terminator elements slowing down the elongating RNAPII to promote termination^16^. Transcription can also be influenced by distal regulatory mechanisms and 3D chromatin conformation^3,17^. Several studies have assessed this readthrough transcription under homeostatic conditions and about 34% of genes have been estimated to produce readthrough transcripts >5kb past the transcription end site (TES)^18–20^.

Cancer cells tend to have more extensive amounts of readthrough transcripts and the amount of readthrough transcripts correlates with poorer survival^21^. Some consequences of readthrough transcription are the formation of RNA chimeras from trans splicing of RNA generated from upstream and downstream genes but these transcripts seem to fail to be translated^22^. Furthermore, formation of antisense RNA can suppress expression of downstream convergently transcribed genes^19,21^.

Both the length and extent of readthrough transcripts are increased following exposure to several different cell stressors such as osmotic stress^23–26^, heat shock^22,24,27^ oxidative stress^24^ and herpes simplex viral infection^22,28–30^. What these stressors seem to have in common is that they interfere with the ability of RNAPII to slow down at the 3’-end of genes and interfere with the activation of the CSF complex resulting in readthrough transcription without cleavage of the 3’-end of the nascent transcript^24,25,31^.

Readthrough transcripts or the act of readthrough transcription may have important functions for the expression of proximal genes^2,32^. Thus, further study of their expression patterns and functions could advance our understanding of transcriptional regulation. Utilizing transcriptomics and functional genomics data generated during the fourth phase of the ENCODE project (ENCODE4), we used nascent RNA Bru-seq to assess the prevalence of transcription in the genomes of 16 human cell lines as well as readthrough transcripts of protein-coding genes under non-stressed conditions. We describe the distinct patterns of readthrough transcripts relative to their associated genes, in addition to identifying sequence and chromatin features that correlate with their transcription. We also used Bru-seq to validate earlier studies showing increased amounts of readthrough transcripts following heat shock, hyperosmotic stress and exposure to arsenic. To further elucidate mechanisms regulating termination of transcription, we treated human cells with drugs targeting histone acetylation or RNAPII phosphorylation as well as used auxin-induced degradation of specific factors known to regulate transcription and assessed their roles in regulating the amounts of readthrough transcripts. These studies extend our understanding of the mechanisms regulating the amounts of readthrough transcripts and identify several factors involved in their regulation.

## RESULTS

### Exploring the transcriptional landscape with the ENCODE4 deeply profiled cell lines

For this study, we used data generated for the ENCODE4 deeply profiled cell lines. This is a unique sample collection composed of data from 16 human cell lines (2 biological replicates, 32 total samples) that were grown in our laboratory to minimize batch effects. These cell samples were then distributed to 8 ENCODE consortium laboratories to perform 13 diverse biochemical assessments (Fig. 1a). Represented in this collection are 6 assays with 9 unique modalities probing different features of transcription, including the steady-state transcriptome (total RNA-seq), RNA synthesis and turnover (Bru-seq and BruChase-seq 2h/6h, respectively)^33^, transcript isoform diversity (long-read RNA-seq), transcription initiation at promoters and enhancer elements (BruUV-seq, PRO-cap), and small non-coding RNA expression (miRNA-seq), as well as the transcriptional response to a perturbation (Bru-seq with ionizing radiation). Additionally, there are 5 assays with 6 unique modalities describing the chromatin landscape of the cell, including accessibility (ATAC-seq, snATAC-seq, DNase-seq) and 3D organization (Intact Hi-C, POLR2A ChIA-PET, CTCF ChIA-PET). Together, these bioassays provide a unique resource to explore the form and function of the human genome in a set of matched biosamples.

**Figure 1.**
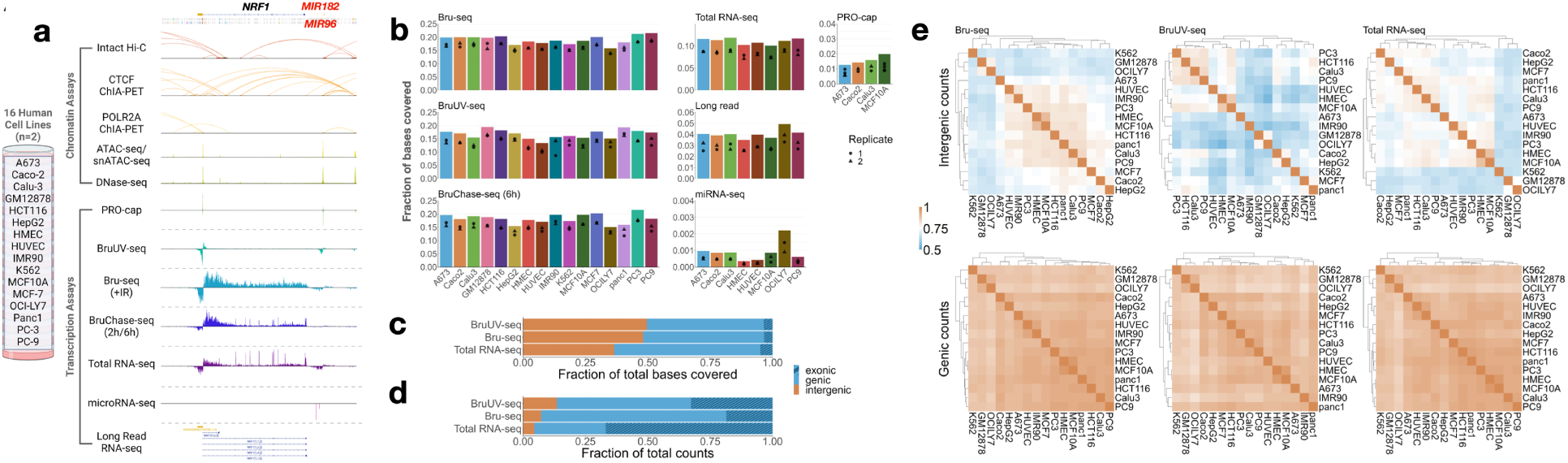
Deeply profiled ENCODE cell lines and genome coverage. **a** Schematic describing the ENCODE4 deeply profiled cell line collection of data (N = 2 biological replicates). Signal tracks are included for a 274.5 kb region containing the NRF gene, MIR182, and MIR96. **b** Genome coverage of sequencing reads for each of the different sequencing assays performed across the cell lines. Cell line coverages are calculated from merged replicate data, thus cell lines with 2 replicate samples per assay are shown. Individual replicate coverages are indicated by the points within the bars. **c** Distribution of bases covered for selected assays (16 cell lines, 24 libraries). The fraction of bases covered was calculated for cumulative genic and intergenic compartments of the genome (based on GENCODE v29), with the striped bar indicating the proportion of genic coverage over exons. **d** total uniquely mapping reads was calculated for cell line specific genic and intergenic compartments of the genome. **e** Intergenic versus genic counts correlations (Pearson’s r) and hierarchical clustering of cell lines are shown for select assays. Color scale shows a shift from moderate (0.5) to high (1) correlations between cell lines, highlighting increased variability in the intergenic space.

To explore the transcriptional landscape, we examined the RNA coverage of the GRCh38/hg38 reference genome using the nascent RNA Bru-seq technique and calculated the cell line specific and cumulative proportion of bases covered on both strands (Fig. 1a) (Supplementary Dataset 1). 18-22% of bases were covered by Bru-seq reads per cell line (Fig. 1b) with 47.6% of the genome covered cumulatively by all cell lines (Supplementary Fig. 1). With RNA-seq (total RNA) data we observe a cumulative genome coverage of about 30%, while long read (poly(A) captured), PRO-cap and miRNA show a cumulative coverage of about 15%, 5% and 1%, respectively (Supplementary Fig. 1).

While these estimates describe the processivity of RNAPII, they do not accurately represent the distribution of transcriptional activity. Thus, we contrasted the coverage fractions and read depth in genic and intergenic compartments and found that ∼50% of all bases covered by Bru-seq data are in the unannotated portion of the genome (Fig. 1c) however, the fraction of reads is predictably higher in genes (Fig. 1d). Despite the smaller fraction of RNA sequencing reads, a significant proportion of genomic space traversed by RNAPII is between genes, and this intergenic activity is more variable among cell lines (Fig. 1e).

### Readthrough transcription is found downstream of most expressed genes

Termination of RNAPII can occur beyond transcription end sites (TESs), resulting in readthrough transcripts. Readthrough transcripts can be detected in the homeostatic total RNA pool^20,32^, and can be further enhanced in response to various stress conditions^24,34^. However, nascent readthrough transcripts making up a significant portion of the transcription occurring in intergenic space have not been comprehensively characterized genome-wide across multiple human cell lines. We implemented an algorithm based on hidden Markov model (HMM) segmentation of Bru-seq data to identify regions of nascent homeostatic readthrough transcription downstream of genes. Briefly, HMM segmentation^35–37^ was performed using 250 bp binned Bru-seq RPKM values as an input, where each segment was assigned an HMM state describing the level of transcription reads in the region. Consecutive segments with the same or decreasing transcription read states were then merged to define a readthrough segment (RTseg) that was assigned to the most upstream gene TES (gene length > 1 kb) it overlapped. RTsegs were defined for each biological replicate Bru-seq sample across the 16 cell lines (Supplementary Dataset 2). We found that readthrough transcription shows a very similar pattern across the 16 cell lines (Fig. 2a). Furthermore, the average distances of readthrough transcripts past the TES (RTsegs) are very similar across the 16 cell lines with an average of 17.4 kb (Fig. 2b). Expressed protein coding genes account for the vast majority of the readthrough reads observed with ∼77% of these genes producing nascent readthrough transcripts (Fig. 2c). Thus, readthrough transcription is a wide-spread phenomenon across human cells.

**Figure 2.**
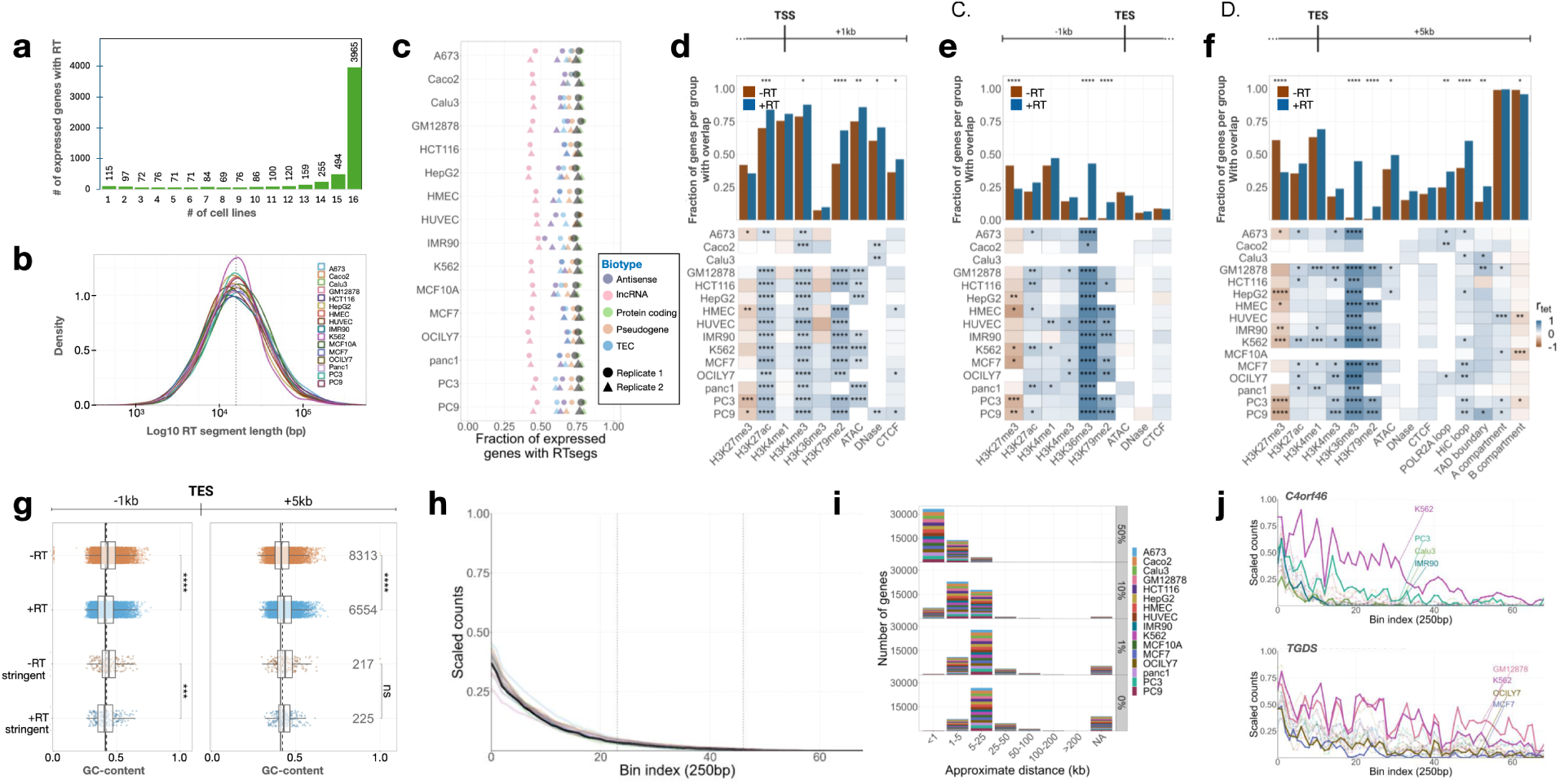
RT transcription is detected far downstream of most expressed genes. **a** The frequency of expressed genes (RPKM > 0.25 in all cell lines, N = 5910) with a readthrough segment (RTseg) in 1-16 cell lines. **b** Log10 length distributions of all RTsegs across the 16 cell lines, where the dashed line indicates the median of each distribution. **c** Fraction of expressed genes per cell line with an assigned RTseg. For both replicates, the fraction of all expressed genes (black) and the fraction of expressed genes broken down by biotype is shown. **d-f** Overlap of stringent readthrough/non-readthrough (RT/ NRT) genes with various chromatin features in the 1 kb downstream **d,** the 1 kb upstream **e**, and 5 kb downstream **f** of the TES. The fraction of RT/NRT genes that had each of these features in any cell line is shown (top), along with the tetrachoric correlation (rtet) between RT/NRT genes and the presence or absence of each feature per cell line (bottom). Fisher’s exact p-values are designated where significant: *p < 0.05, **p < 0.01, ***p < 0.001, ****p < 0.0001. **g** The distribution of GC-content for all and stringent RT and NRT genes for the sequences 1 kb upstream (left) and 5 kb downstream (right) of gene TESs. Vertical lines represent the genome-wide GC-content15 (solid), and the median GC-content of all 1 kb or 5 kb regions (dashed). FDR adjusted p-values were determined from Wilcoxon tests are denoted as: *padj < 0.05, ** padj < 0.01, *** padj < 0.001, **** padj < 0.0001. **h** The median aggregated scaled signal of all 17.3 kb RT signals (black) and cell line specific RT signals. Scaled counts are calculated for 250 bp bins (Bin index) and vertical lines demarcate 23-bin sectors used downstream of the TES. **I**) Distributions of genes with RT signal that reaches 0-50% of the 3’-gene signal within the approximate distance ranges shown. NA indicates the number of genes with RT signals that never reach the threshold, and the color shows the even distribution of genes per cell line. **j** Scaled RT signals downstream of the C4orf46 and TGDS genes are shown for all cell lines. Selected cell lines are highlighted to display interesting patterns of variability.

Comparing genes displaying readthrough transcripts with genes lacking it we found that genes showing readthrough transcripts showed enrichment in the beginning of their genes in histone modifications such as H3K27ac, H3K4me3, H3K79Kme2. Furthermore, genes with readthrough transcripts showed a more open chromatin structure as determined with ATAC-seq (Fig. 2d). At the end of and downstream of the genes we observed that the predominant histone modifications associating with readthrough transcripts were H3K36me3 and H3K79me2 (Fig. 2e&f). Thus, genes with readthrough transcription have distinct histone marks associated with them at the beginnings, ends and downstream of these genes. We next examined whether the GC content in the DNA 1 kb upstream and 5 kb downstream of the TES affects readthrough transcription. It was found that readthrough genes tend to have a lower GC-content at the end of genes while having a higher GC-content downstream of the TES (Fig. 2g).

### Readthrough transcription varies among genes and cell lines

Rather than terminate in a defined manner, readthrough signal tends to taper off gradually^14,24,38,39^. To better understand this behavior, we calculated binned counts (250 bp) for readthrough transcripts and scaled these to the signal at the end of the gene (∼1 kb upstream of the TES) (Fig. 2h). With these scaled counts, we assessed the approximate distance required for each readthrough signal to reach 50, 10, 1, and 0% of the signal at the 3’-end of the gene in the 16 cell lines (Fig. 2i). We found the readthrough signal for most genes was reduced below 50% within the first 1 kb downstream of the TES. The distances the readthrough transcripts reach past the TES show an exponential decay and most genes show a decrease to 0 scaled counts within 25 kb. However, there is great variability between genes with a wide range of signal and lengths observed. Furthermore, we found variability between cell lines as well (Fig. 2j). For example, for the *C4orf46* gene there are higher levels of readthrough transcripts in K562 and PC3 cells than for all the other cell lines. Likewise, in the *TGDS* gene we found strong and extended readthrough in the GM12878 and K563 cells compared to the other cell lines. Thus, readthrough transcription shows both gene and cell line-specific behaviors.

### Cell perturbations causing increased readthrough transcripts

Exposure of cells to heat shock, oxidative stress or hyperosmotic stress have been shown to result in enhanced readthrough transcription as assessed by various RNA assays^22–27^. In addition to causing readthrough transcription, these stressors also rapidly induce molecular condensate formation in cells^26,40^. It has been suggested that the sequestering of cleavage and polyadenylation factor 6 (CPAF6) to these condensates may prohibit transcription termination at the end of genes during these stress conditions^26^. Here we used nascent RNA Bru-seq to assess readthrough transcription in U2OS osteosarcoma cells exposed to heat shock, sodium arsenite (oxidative stress) or hyperosmotic stress. To compare the effects of exposure to different stressors with mock-treated controls, we scaled the reads by calculating binned counts (50 bp) for readthrough transcripts and compared them with the signal at the end of the gene (∼1 kb upstream of the TES).

Following 60 minutes of heat shock (42°C) of U2OS cells, we observed 904 genes transcriptionally downregulated and 1363 genes upregulated (>2-fold, padj<0.01). (Supplementary Fig. 2a, Supplementary Dataset 3). Performing gene set enrichment analysis (GSEA) we find strong upregulation of “response to heat” and “HSF1 dependent transcription” (Supplementary Fig. 2b). Among the up-regulated genes we find heat shock genes such as *HSPA6* and *HSPH1* (Supplementary Fig. 2c). These findings indicate that a robust heat shock response was elicited in these cells after a 60-minute incubation at 42°C. In a small set of genes, such as *RAB3IP*, *HIKESHI* and *MRPS6*, we observed increased amounts of readthrough transcripts after heat shock (Supplementary Fig. 2c). To assess whether heat shock promotes nascent readthrough transcription genome-wide, we compared the transcription reads downstream of TES in control cells (37°C) with transcription reads in heat shocked cells (42°C, 60 min) and scaled these reads with reads from transcription from 1 kb upstream of the TES. Comparing these scaled reads, we observe only a slight genome-wide increase in nascent readthrough transcripts following heat shock compared to control cells (Fig. 3a).

**Figure 3.**
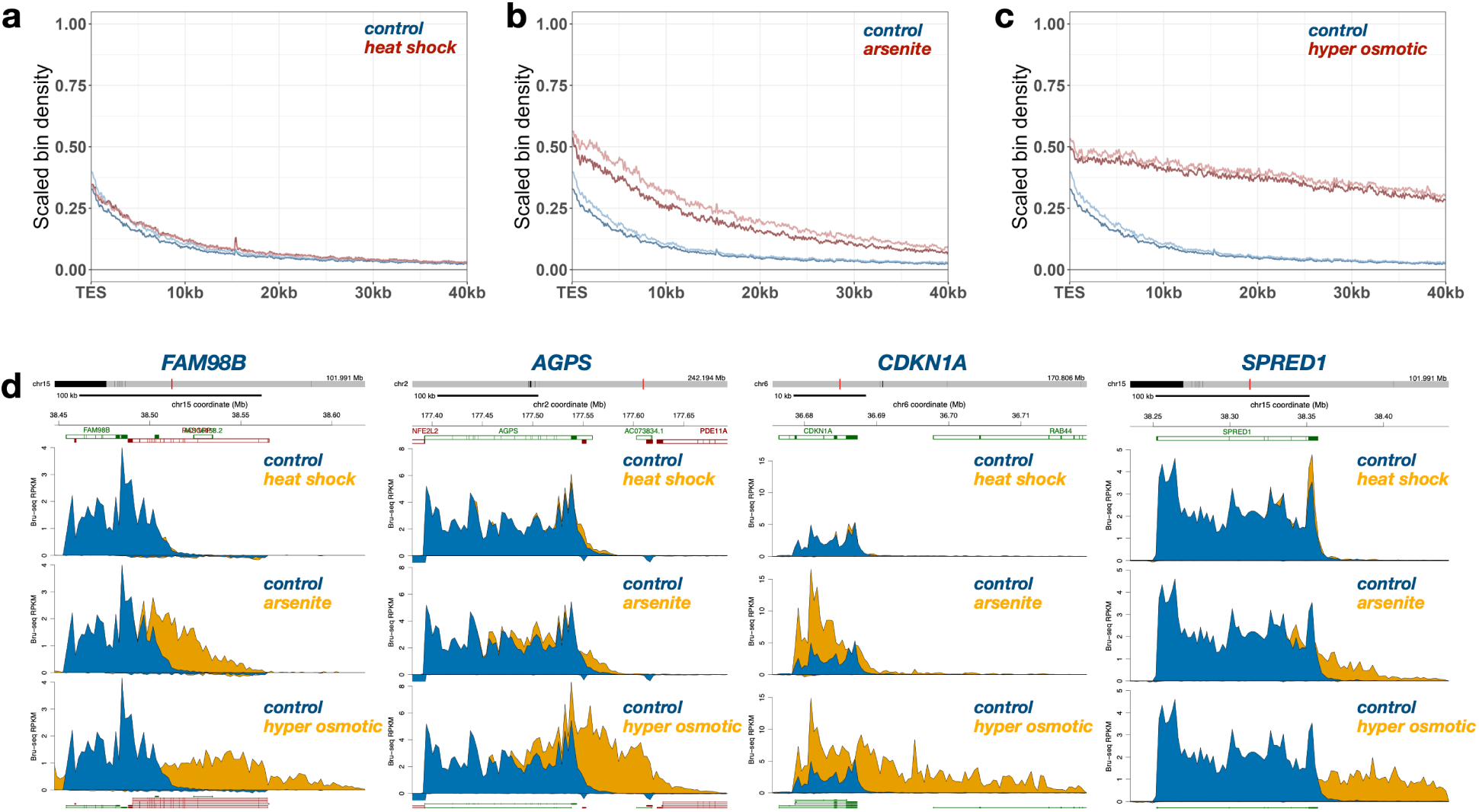
Cellular stress causes an increase in readthrough transcripts. **a** Scaled readthrough transcripts comparing control U2OS cells and heat shock-treated cells (1-hour incubation at 42°C). **b** Scaled readthrough transcripts comparing control U2OS cells and cells treated with sodium arsenite (0.5 mM for 1 h). **c** Scaled readthrough transcripts comparing control U2OS cells with cells treated with hyperosmotic stress (300 mM sodium, for 1 h). **d** Bru-seq browser images of four genes where reads from control cells are in blue and from treated cells in yellow.

We next treated the U2OS cells with sodium arsenite (0.5 mM, 60 min) which is known to induce oxidative stress^41,42^. Using nascent RNA Bru-seq, we found that 3347 genes were downregulated and 2020 genes were transcriptionally upregulated following treatment (Supplementary Fig. 3a, Supplementary Dataset 4). Gene sets upregulated include “coagulation” and “HSF1 dependent transactivation” while “oxidative phosphorylation” and “DNA repair” sets were downregulated (Supplementary Fig. 3b). We found that many genes upregulated by heat shock were also induced after arsenic treatment such as *HSPH1* (Supplementary Fig. 3c). Several genes, such as *NEDD1*, *RELCH*, *STAM* and *KDM5A*, showed strong increase in readthrough transcripts following exposure to arsenite (Supplementary Fig. 3d). When comparing the genome-wide aggregate of scaled reads, we observe that arsenite treatment results in a robust increase in readthrough transcripts (Fig. 3b). The increase in readthrough transcript signal was clearly evident immediately past the TES suggesting that arsenite suppresses the cleavage of the 3’ends of nascent transcripts. No obvious change in the shape of the curve downstream of the TES compared to control cells was observed indicating no observable effect of arsenite treatment on the normal mechanism of transcription termination.

Hyperosmotic stress (300 mM NaCl for 60 min) resulted in a dramatic change in transcription with 5547 genes downregulated and 6077 genes upregulated (>2-fold, padj<0.01) (Supplementary Fig. 4a, Supplementary Dataset 5). Some gene sets enriched included “KRAS signaling down” and “coagulation” and downregulation of gene sets “E2F targets” and “DNA repair” (Supplementary Fig. 4b). Many genes showed evidence of readthrough transcription following hyperosmotic stress such as *STAM, ATF1, AGPS, LIMCH1, and XRCC5* (Supplementary Fig. 4c). When aggregating the scaled reads, we observed that hyperosmotic stress induces a dramatic increase in readthrough reads and this increase can be seen immediately past the TES and also the reads show a much slower decay slope (Fig. 3c). These results suggest that hyperosmotic stress interferes with both the 3’-end cleavage of nascent transcripts and in the rate of termination of transcription of the RNAPII elongating past the TES.

Comparing readthrough transcription of the three different treatments across genes we see that heat shock has no observable effect of readthrough transcription of any of the four gene examples (Fig. 3d) while we did observe readthrough of a small set of genes (Supplementary Fig. 2d). Both arsenite and hyperosmotic stress caused an increase in readthrough reads for these four genes which were most severe for the samples treated with hyperosmotic stress. Arsenite and hyperosmotic stress, but not heat shock, induced transcription of the p53-inducible gene CDKN1A suggesting that these stresses activate the p53 response (Fig. 3d).

### HDAC inhibitor Vorinostat does not affect readthrough transcription

To help elucidate the mechanisms responsible for regulating readthrough transcription, we next explored whether the histone deacetylase (HDAC) inhibitor Vorinostat affects readthrough transcription. We used the sarcoma cell line HOS and treated them with 10 µM Vorinostat and assessed nascent transcription during the last 30 minutes of a 16h treatment. Vorinostat is known to induce histone hyperacetylation, chromatin decondensation and increased transcription^43,44^. The hypothesis to be tested was that histone hyperacetylation leading to a more open chromatin structure may promote longer readthrough transcription by enhanced elongation rates potentially making it more difficult for XRN2 to catch up with elongating RNAPII past the termination site or promoting a chromatin structure less prone to form roadblocks. The results show that inhibition of histone deacetylases had a large effect on nascent transcription with 3,233 genes downregulated and 3,496 genes upregulated (Supplementary Fig. 5a, Supplementary Dataset 6). GSEA analyses revealed that gene sets such as “UV response up” and “KRAS response down” showed enhanced expression while expression of “MYC targets” and “E2F targets” were suppressed (Supplementary Fig. 5b). Genes such as *STC1* and *AREG* were dramatically induced while *IL7R* and *FLI1* were repressed following HDAC inhibition (Supplementary Fig. 5c). Aggregation of scaled reads downstream of TES showed no discernable difference compared to control cells, suggesting that chromatin remodeling due to histone hyperacetylation by Vorinostat does not affect readthrough transcription (Supplementary Fig. 5d).

### Increased readthrough transcripts following inhibition of splicing

We and others have found that co-transcriptional splicing is inefficient leading to a sizable portion of nascent transcripts remaining fully unspliced^33,45–48^. It has been suggested that transcripts that are not spliced are not cleaved at the Poly(A) site at the 3’-end and are retained attached to chromatin^49,50^. Furthermore, using long read sequencing it has been found that readthrough transcripts are predominantly unspliced^45,46^. Thus, it is possible that inhibition of splicing may increase readthrough transcription due to loss of 3’-end cleavage. To test this hypothesis, we first explored whether the splicing inhibitor meayamycin B, which targets the SF3b splicing complex^51^, affects readthrough transcription. We first validated that meayamycin B blocks splicing and while control K562 cells showed strong enrichment of exonic reads in the *MBNL1* transcript after a 6-hour chase, the meayamycin B treated cells did not show that suggesting that splicing is completely inhibited (Fig. 4a). When analyzing the scaled aggregate plot of readthrough transcripts, inhibition of splicing resulted in an increase of these transcripts (Fig. 4b). The readthrough transcripts are more abundant immediately past the TES with a similar profile of decay away from the TES. We next explored the rate of turnover of the readthrough transcripts by chasing the Bru-labelled cells for 2 hours (Fig. 4c) or 6 hours (Fig. 4d). It is observed that readthrough transcripts in control cells were much more rapidly turned over than the readthrough transcripts in the Meayamycin B treated cells. Examples of 4 genes that show strong enhancement of readthrough transcripts following splicing inhibition are shown in Figure 4e (Supplementary Dataset 7).

**Figure 4.**
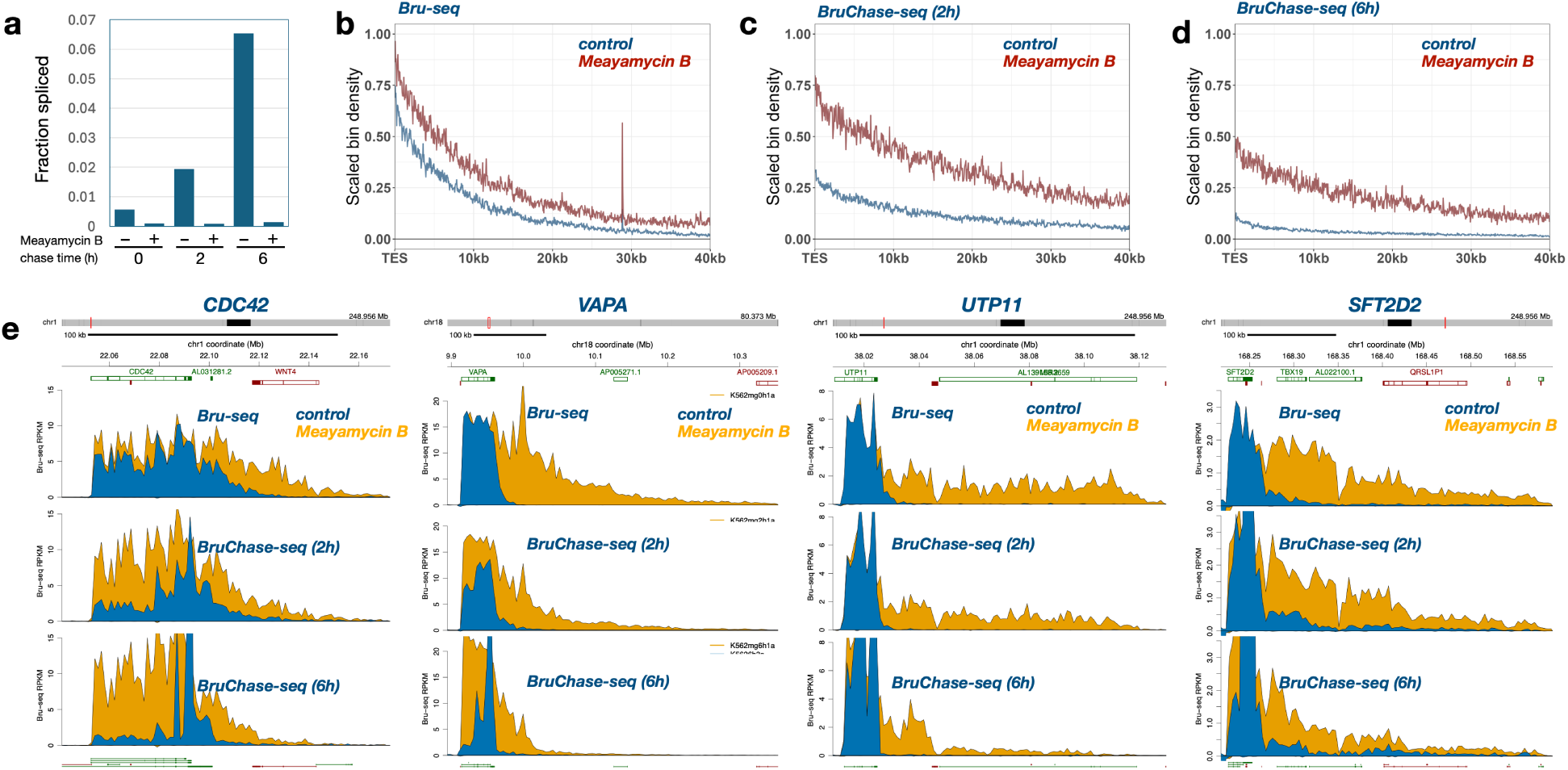
Inhibition of splicing leads to enhanced readthrough transcripts. **a** K562 cells were either mock treated or treated with Meayamycin B (3 µM for 6 h at 37°C) and Bru-labeled RNA was isolated and sequenced either directly after Bru-labeling (0 h) or after a 2 or 6 hour chase. The fraction of spliced reads with and without Meayamycin are plotted. Scaled readthrough transcripts were plotted and compared between control and Meayamycin B treated K562 cells directly after Bru-labeling **b** after a 2 h chase **c** or a 6 h chase **d. e** Bru-seq browser images of four genes where reads from control cells are in blue and from treated cells in yellow.

We next used another SF3B inhibitor, pladienolide B^52^ (plad B) to validate the results we obtained with Meayamycin B. Normal human fibroblasts were treated for 1 hour with 100 nM plad B and transcriptional effects were analyzed immediately using Bru-seq or after a 6-hour chase using BruChase-seq. Plad B was found to have a significant effect on splicing when comparing the “fraction spliced” after a 6-hour chase (Supplementary Fig. 6a). We found that 402 genes were downregulated and 109 genes upregulated (>2-fold, padj<0.01) when analyzed with Bru-seq (Supplementary Fig. 6b, Supplementary Dataset 8). GSEA analysis shows that “UV response DN” and “mitotic spindle” were enriched and “TNFa signaling via NFKb” and “E2F targets” were suppressed (Supplementary Fig. 6c). When exploring readthrough transcripts genome-wide using the scaled aggregate analysis we found that there was an increase after plad B treatment (Supplementary Fig. 6d). We observed several genes with increased readthrough transcripts such as *TLR4, UBE2E3, ARIH1* and *CDC73* (Supplementary Fig. 6e).

### Auxin-mediated degradation of XRN2 leads to increased readthrough transcripts

The 5’-3’ RNA exonuclease XRN2 is known to participate in the degradation of readthrough RNA formed downstream of the TES after cleavage by the 3’-end cleavage complex and is the main enzyme responsible for transcription termination through the torpedo mechanism^10–12,53^. The XRN2 acts very rapidly on cleaved pre-mRNA so this RNA can only be seen in cells depleted of XRN2^13^. To test whether the absence of XRN2 would cause an increase in readthrough signal as determined by nascent RNA Bru-seq, we used HCT116 cells with auxin-inducible degradation of the XRN2 protein^54^. As expected, it was found that depletion of XRN2 resulted in a dramatically enhanced level of readthrough signal immediately after the TES (Fig. 5a). This increased readthrough signal most likely stems from transcripts that were cleaved at the 3’-end but the rapid XRN2-mediated transcription termination is not taking place^13^. However, the readthrough read decay is very similar to the decay rate in control cells suggesting that the termination of 3’-end cut and uncut nascent transcripts are subjected to the same rules of transcription termination. The expression of XRN2 in the auxin responsive cells is much lower even in the absence of auxin than levels found in parental cells (Supplementary Fig. 7) which explains the higher level of readthrough transcripts in the control auxin responsive than we see in the parental cells. Zooming in on individual genes, Bru-seq data show increased readthrough transcript levels following XRN2 knockdown in several genes such as *TGIF1, SMAD5, VEGFA, ETNK1* and *ACADM* (Fig. 5b, Supplementary Dataset 9).

**Figure 5.**
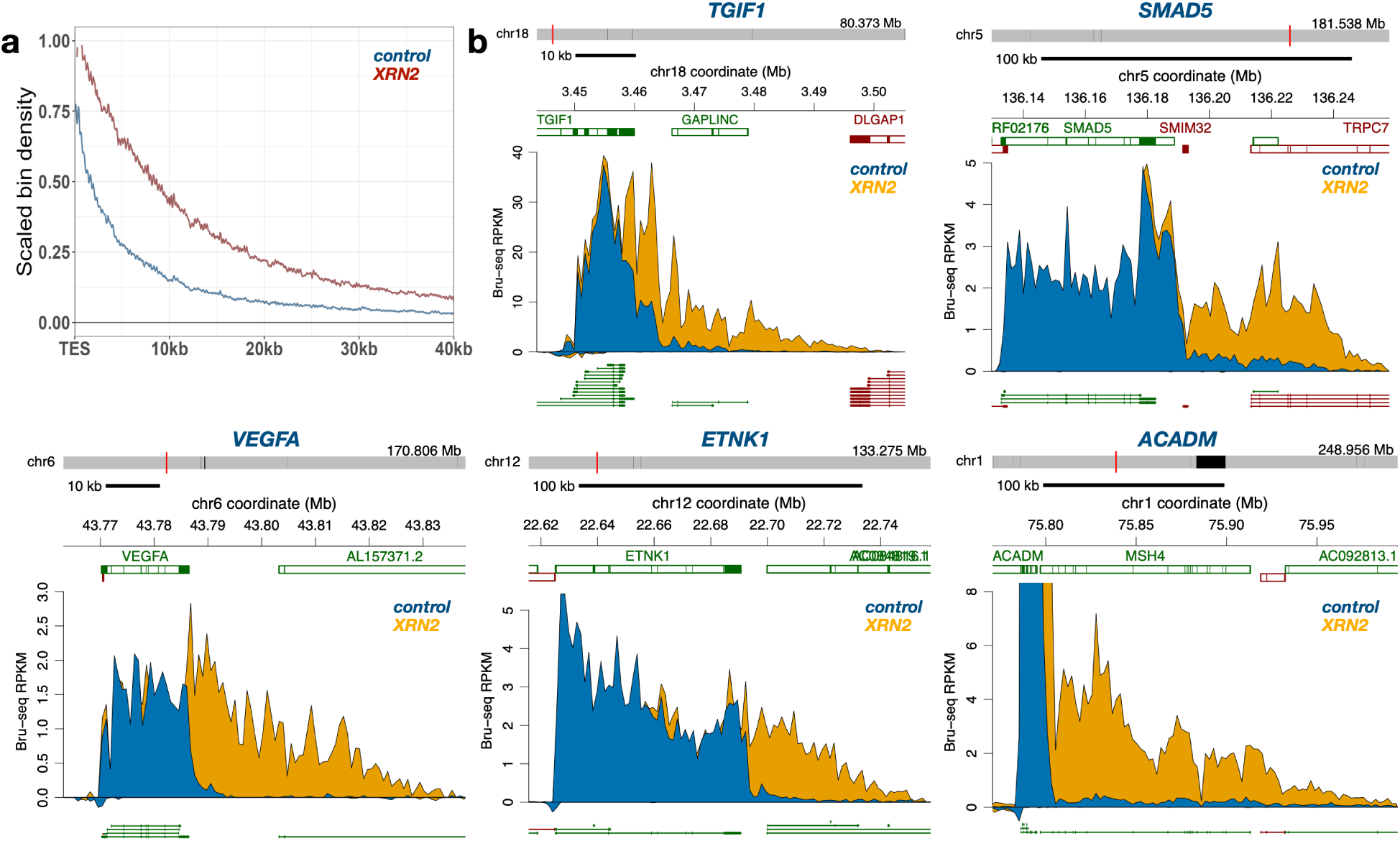
Loss of XRN2 leads to enhanced levels of readthrough transcripts. **a** Scaled readthrough transcripts were plotted and compared between control HCT116 and HCT116 cells treated with auxin to degrade XRN2. **b** Bru-seq browser images of five genes where reads from control cells are in blue and reads from XRN2-depleted cells in yellow.

### No role observed for RNA exosome, the cohesin complex, SMARCA5 or CDK7 in regulating readthrough transcripts

It has been suggested that the RNA exosome participates in the termination of transcription in yeast^55,56^. However, no effect on readthrough transcription as measured by nuclear RNA-seq was observed in cells depleted of either DIS3 or EXOSC10 subunits of the RNA exosome^54^.

Here we show that nascent RNA Bru-seq concurs with the data from nuclear RNA-seq^54^ in that no significant effects on the readthrough signal were observed after DIS3 or EXOSC10 depletion in HCT116 cells (Supplementary Fig. 8a&b, 9, 10, Supplementary Datasets 10, 11).

We next explored how loss of either of two chromatin domain organizing proteins, RAD21 and CTCF would affect readthrough transcription. CTCF is an architectural protein which forms chromatin loop domains, or topological associated domains (TADs), through ATP-dependent loop extrusion by the cohesin complex^57–60^ while Rad21 is a subunit of the cohesin complex^61^. Treatments with auxin caused dramatic depletion of the proteins already after 30 minutes^54^. Examining readthrough transcripts in HCT116 cells depleted of either CTCF or RAD21 showed no apparent difference compared with control cells (Supplementary Fig. 8c&d, Supplementary Datasets 12, 13).

We next explored whether depletion of the ATPase chromatin remodeler SMARCA5 (SNF2H) which is critical for the maintenance of proper spacing of nucleosomes in the genome^62,63^, may affect readthrough transcription. SMARCA5 has been shown to interact with both CTCF and RAD21 and depletion of SMARCA5 leads to the remodeling of chromatin in and around CTCF binding sites leading to the loss of CTCF binding to chromatin^63^. Here we show that depletion of this chromatin remodeler^63^ has no noticeable effect on the level of readthrough transcripts (Supplementary Fig. 8e, Supplementary Dataset 14). We also depleted CDK7 in HCT116 cells and performed Bru-seq analysis. CDK7 is the catalytic component of the CDK activating kinase (CAK) and phosphorylates the C-terminal domain (CTD) of RNA Polymerase II to^64^. Here we found that depletion of CDK7 did not affect readthrough transcription (Supplementary Fig. 8f, Supplementary Dataset 15).

### Role of MED14, BRD14 and SUPT16H in regulating readthrough transcripts

MED14 is the largest subunit of the Mediator complex that serves as a scaffold connecting regulatory proteins to the basal transcription machinery^65^. The Mediator complex also coordinates RNA polymerase II transcription elongation, splicing and termination^66^. Depletion of MED14 in HCT116 cells resulted in an increase in readthrough transcripts (Fig. 6a). Depletion of MED14 resulted in 1960 genes transcriptionally downregulated and 1990 genes upregulated (Supplementary Fig. 11a, Supplementary Dataset 16). GSEA analysis revealed upregulation of pathways such as “interferon gamma response” and “xenobiotic metabolism” while downregulation of “MYC targets” and “ribosome” (Supplementary Fig. 11b). Some genes dramatically downregulated are *MYC* and *WNT16 and* several genes, such as *PHF20L1*, *PDLIM5* and *SSBP1* showed an increase in readthrough transcripts following MED14 depletion (Supplementary Fig. 11c&d).

**Figure 6.**
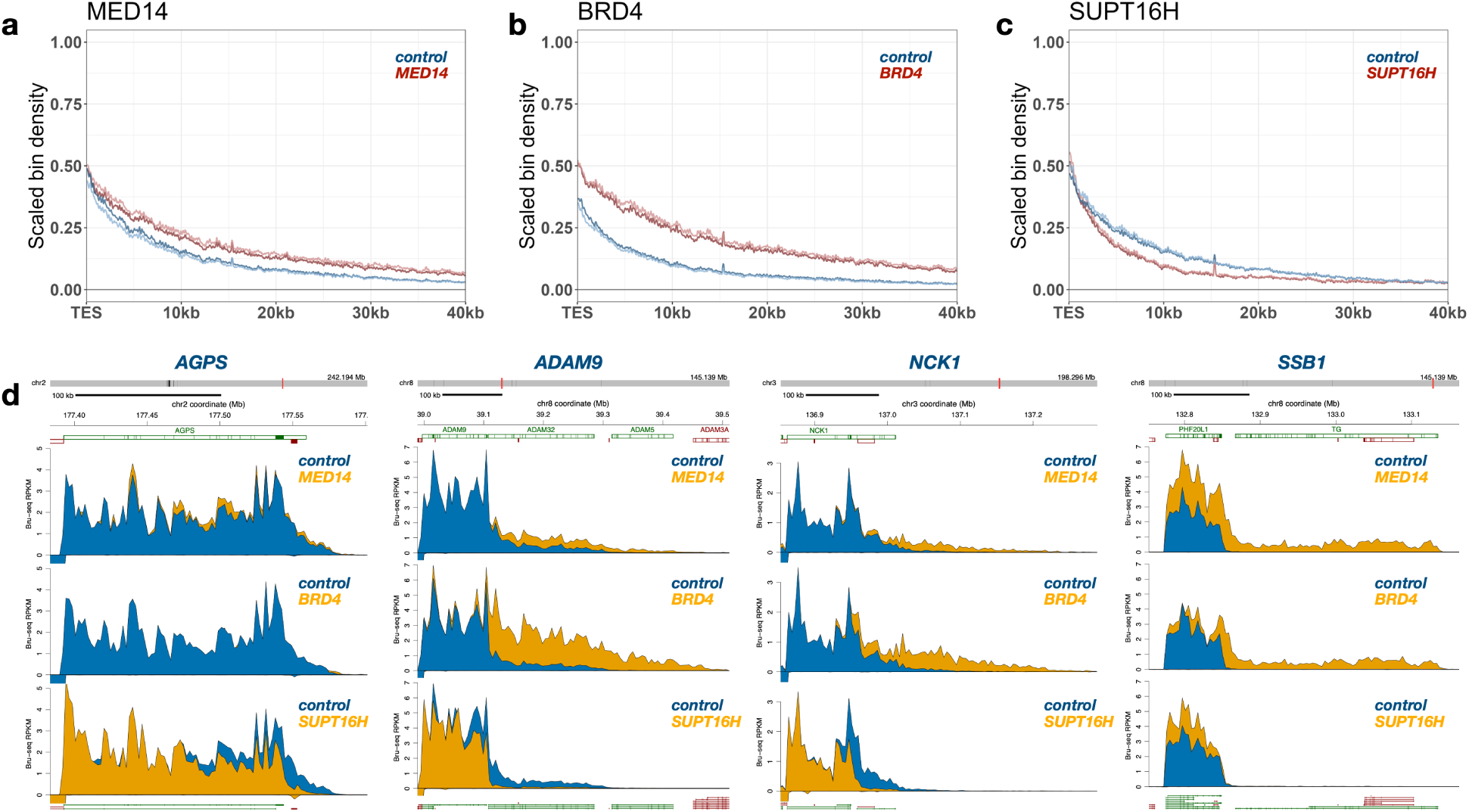
Depletion of MED14 or BRD4 increases while depletion of SUPT16H decreases levels of readthrough transcripts. **a** Scaled readthrough transcripts were plotted and compared between mock (blue) and auxin-treated (red) HCT116 cells harboring a MED14 gene carrying an AID tag for auxin-mediated degradation. **b** Scaled readthrough transcripts were plotted and compared between mock (blue) and auxin-treated (red) HCT116 cells harboring a BRD4 gene carrying an AID tag for auxin-mediated degradation. **c** Scaled readthrough transcripts were plotted and compared between mock (blue) and auxin-treated (red) HCT116 cells harboring a SUPT16H gene carrying an AID tag for auxin-mediated degradation. **d** Bru-seq browser images of four genes where reads from control cells are in blue and from auxin-treated cells in yellow.

BRD4 is a bromodomain protein that binds to acetylated histones^67^ and regulates transcription pause release by controlling the positive elongation factor P-TEFb^68^. There is also evidence that BRD4 plays a role regulating 3’-processing and restricting readthrough transcription^69^. To further test the role of BRD4 regulating readthrough transcription we depleted HCT116 cells of BRD4 and performed Bru-seq. A clear increase in readthrough transcription was observed in the absence of BRD4 validating its role in confining readthrough transcription (Fig. 6b). Depletion of BRD4 resulted in significant changes in the nascent transcriptome with 1892 genes downregulated and 2555 genes upregulated (Supplementary Fig. 12a, Supplementary Dataset 17). “Spermatogenesis” is a pathway that was upregulated while “E2F targets”, “estrogen response early” and “DNA replication” were suppressed (Supplementary Fig. 12b). Zooming in on individual genes we found that *HAS3* and *WNT16* were downregulated after BRD4 depletion and several genes showed increased readthrough transcripts such as ADAM9, ACADM and PDLIM5 show dramatic increases in readthrough transcripts (Supplementary Fig. 12c&d).

Another factor critical for chromatin remodeling is SUPT16H which is a subunit of the transcription elongation factor FACT required for transcription on chromatin templates^70,71^. SUPT16H acts as a histone chaperone that promotes the redeposition of H3-H4 histones displaced by the elongating RNA polymerase^72^. Depletion of SUPT16H results in reduced elongation rates of the RNA polymerase and activation of alternative polyadenylation sites^73^. Our results show that depletion of SUPT16H in HCT116 cells leads to a reduction of readthrough transcripts (Fig. 6c). Furthermore, depletion had a major effect on nascent transcription with 2315 genes downregulated and 2544 genes upregulated following SPT16H depletion (Supplementary Fig. 13a, Supplementary Dataset 18). GSEA pathways upregulated included “heme metabolism” and “interferon alpha response” while “MYC targets” and “spliceosome” were downregulated (Supplementary Fig. 13b). Genes downregulated in the absence of SUPT16H included *EGFR* and *MIR17HG* while *FOS* was upregulated and *CDC73* and *NCKAP1* showed reduced amounts of readthrough transcripts because of depletion of SUPT16H (Supplementary Fig. 13c). Thus, the FACT complex plays a role in promoting readthrough transcripts.

## DISCUSSION

While much remains to be discovered regarding the regulation and function of readthrough transcripts, several features of readthrough transcripts and their parent genes have been identified. Readthrough transcripts are typically long, with transcription downstream of genes detected anywhere from a few hundred bases to several hundred kb^23,32^. Genes that give rise to readthrough transcripts are commonly enriched for elongation-related histone marks, such as H3K36me3, at their 3’-ends and downstream of their TES. Furthermore, they are often depleted for GC-rich sequences and have low GC-content at their 3’-ends^20,24,34^. One possible function of readthrough transcripts, or the act of their transcription, is that they are important for the regulation of nearby genes, perhaps through maintenance of open chromatin^24^. A previous study suggested a more specific role for readthrough transcription in reducing mRNA translation of the parent gene, IRF1, in response to viral infection, thus possibly functioning to dampen the host immune response ^30^. As our understanding of these transcripts and the mechanisms underlying their transcription increases, it will be important to uncover why some readthrough transcripts are upregulated under various stress conditions and what role these RNAs may play in normal cellular function.

In this study, we interrogated readthrough transcripts genome-wide across 16 human cell lines profiled with nascent RNA Bru-seq as part of the ENCODE 4 project. Calculating the percentage of bases in the genome (hg38) covered by Bru-seq reads we observe that 18-22% of bases were covered by nascent Bru-seq reads per cell line (Fig. 1). Calculating the cumulative coverage when considering reads from all 16 cell lines we observe that about 48% of the genome show evidence of transcription as compared to about 40% of the genome consisting of protein-coding genes https://www.gencodegenes.org/human/release_29.html (Supplementary Fig. 1). When analyzing the coverage of reads from RNA-seq data from the same cell lines we observe a cumulative genome coverage of about 30%. Previous estimates of transcription coverage of the human genome were higher with about 39% found to be covered by primary transcripts per cell line and with cumulative transcription across 15 cell lines reaching about 75%^74^. The discrepancy in these numbers may stem from different sequencing methods and read depth used.

Using nascent RNA Bru-seq, readthrough transcripts were detected past the TES of more than 75% of protein-coding genes under normal homeostatic conditions (Fig. 2). These readthrough transcripts averaged ∼17 kb in length and both the genes with readthrough transcripts and their lengths were very consistent across the cell lines. Genomic featured that discriminated genes with readthrough transcripts from genes without were enrichment at the beginning of their genes with histone modifications such as H3K27ac, H3K4me3, H3K79Kme2 and showed a more open chromatin structure as determined by ATAC-seq. Furthermore, H3K36me3 and H3K79me2 histone marks were enriched at gene ends and downstream of their TESs and that they presented with a lower GC-content at the gene ends while having a higher GC-content downstream of the TES. The amount of transcript reads show a ∼50% drop just past the TES which most likely represents rapid XRN2-mediated degradation of the RNA synthesized by RNAPII following 3’-end cleavage of the nascent transcript^13–15^. The remaining readthrough reads most likely represent transcripts that were not cleaved at the 3’ ends^13–15^.

Several cellular stressors have been shown to increase both the extent and length of readthrough transcripts ^22–30^. Here we used Bru-seq to assess nascent readthrough transcripts following exposure of human cells to heat shock, sodium arsenite and hyperosmotic stress (Fig. 3). The extent of increased readthrough transcripts following heat shock was overall not very impressive, however, there were several genes that showed increased readthrough transcripts (Supplementary Fig. 2d). Sodium arsenite and hyperosmotic stress cause a much more dramatic increase in readthrough reads. This increase was observed immediately past the TES suggesting that these stresses reduce the 3’-end cleavage of the nascent transcript leading to readthrough transcripts that are not amenable to XRN2 degradation since they do not have a free 5’-end^24,25,31^. Indeed, we previously showed that the cleavage and polyadenylation factor CPSF6 becomes sequestered in molecular condensates following exposure to hyperosmotic stress consistent with decreased 3-end cleavage and enhanced readthrough transcript^26^. In addition to having an initial higher level of readthrough transcripts, the decay rate of signal when moving away from TESs is clearly lower following exposure to hyperosmotic conditions (Fig. 3c). This suggests that hyperosmotic stress leads to both reduced 3-end cleavage of the nascent transcript and to reduced levels of terminating transcription blocks downstream of the TES.

RNA splicing is a fundamental process to excise introns and stitch together coding exons to form mature mRNAs. Splicing is a complex process executed by the spliceosome co-transcriptionally shortly after the nascent RNA exits the elongating RNA polymerase II^75–78^. The RNA spliceosome consists of the small nuclear ribonucleoprotein particles U1, U2, U4, U5, and U6 snRNPs and more than 100 additional proteins^79^. Surprisingly, many primary transcripts do not appear to be spliced at all and it has been suggested that this splicing-uncoupled transcription may function as a genome scanning mechanism to find and repair DNA damage^80^. Furthermore, there is a link between splicing and 3-end cleavage so that unspliced transcripts will not recruit the 3-end cleavage and polyadenylation complex and should therefore generate readthrough transcripts not susceptible to XRN2 degradation and termination^81^ ^82^ ^83^. Indeed, long read sequencing has revealed that the population of readthrough transcripts is exclusively made up of unspliced transcripts^45,46^. Here we show that suppression of splicing with the inhibitors Meayamycin B or pladienolide B increases the amounts of readthrough transcripts (Fig. 4, Supplementary Fig. 6). This is consistent with the notion that transcripts that are not spliced are also not cleaved leading to increased levels of readthrough transcripts.

To obtain mechanistic insights to the regulation of readthrough transcripts an auxin-inducible degradation system in HCT116 cells was utilized^54^ ^84^ ^85^. As expected, degradation of XRN2 resulted in a significant increase in readthrough reads immediately past the TES since RNA polymerase II with a cleaved transcript will not be subjected to the torpedo process by XRN2 (Fig. 5a). The decay curve of the readthrough transcripts away from the TES was similar with or without XRN2 suggesting that the termination mechanism is the same whether the elongating polymerase is attached to a short RNA cleaved at its 5’-end or if it contains a long unspliced transcript. We then depleted HCT116 cells of a series of proteins using the auxin-inducible degradation system and assessed the effect of the depletion of the individual proteins on the extent of readthrough transcripts. We found no observable effect on readthrough transcripts levels following depletion of the RNA exosome components DIS3 or EXOSC10, the chromatin structure organizing proteins RAD21 and CTCF, the chromatin modulator SMARCA5 or the cyclin-dependent kinase 7 (CDK7) (Supplementary Fig. 8). In contrast, depletion of MED14 and BRD4 resulted in increased readthrough transcripts while degradation of SUPT16H caused the readthrough transcripts to become shorter (Fig. 6)

Our studies document a pervasive readthrough transcription for over 75% of protein-coding genes in human cells during homeostatic conditions. Examining the scaled aggregate plots of readthrough transcripts, about half of the primary transcripts are lost immediately past the TES most likely due to the rapid degradation by XRN2 from the 5’-end of the cleaved RNA still attached to the RNA polymerase II elongating past the TES^13–15^. The other half of the readthrough transcripts that we capture with the Bru-seq technique most likely represents primary transcripts that did not get co-transcriptionally spliced and therefore did not activate the 3-end cleavage process when reaching the TES. This model is consistent with previous studies using long-read RNA sequencing showing that splicing exhibits a binary “all or none” splicing pattern^45–48^. We recently confirmed these findings using Bru-seq and found that about half of the nascent transcripts were co-transcriptionally spliced while the other half appeared to not be spliced at all^33^. Furthermore, inhibition of splicing showed an increase in readthrough transcripts, again linking lack of splicing to the appearance of readthrough transcripts. While the fully spliced transcripts will generate mRNA that can be exported and translated, the splicing uncoupled transcription may be used for genome surveillance to detect and repair DNA damage through transcription-coupled repair^80^ (Fig. 7a).

**Figure 7.**
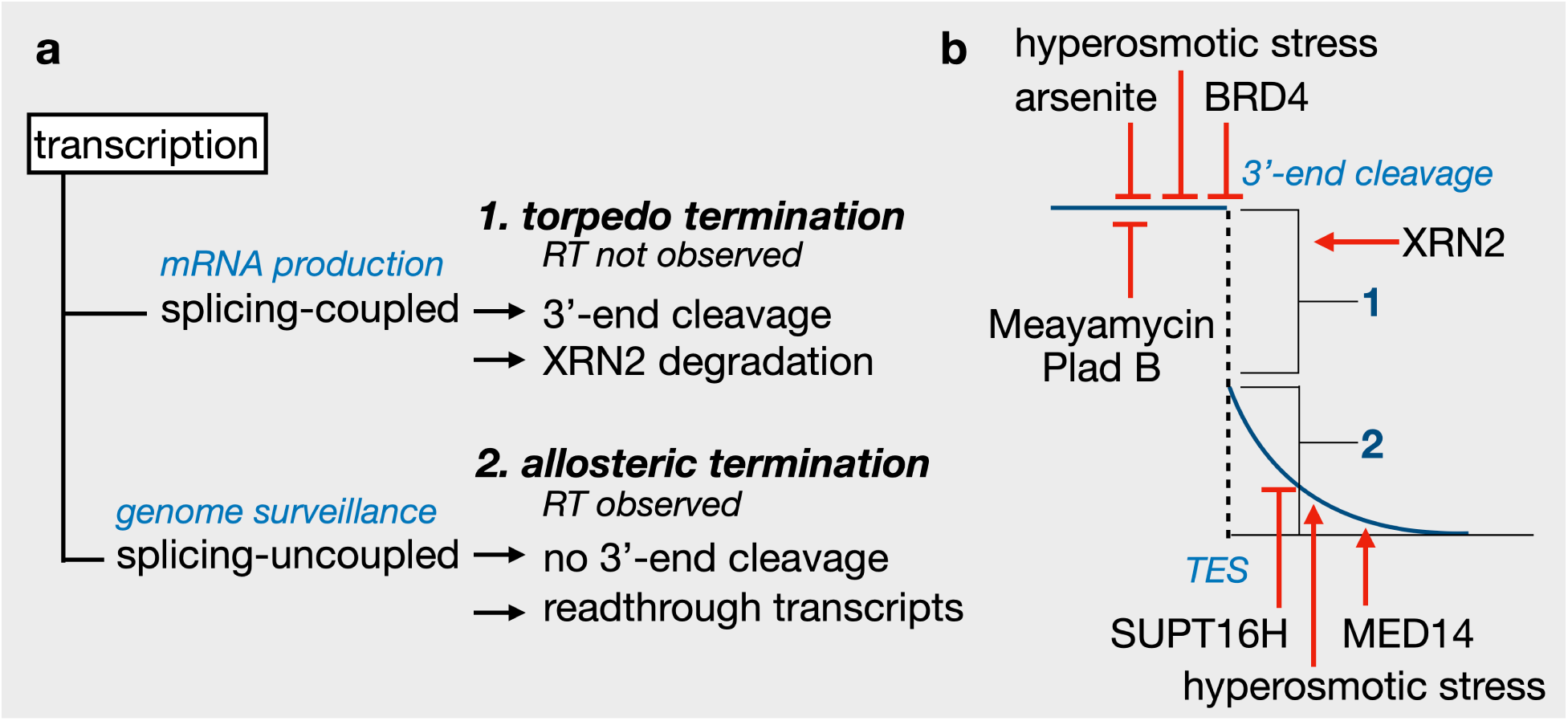
Model for regulation of readthrough transcripts. **a** Transcription is either splicing-coupled or splicing-uncoupled. If transcripts are spliced, they will be processed by the 3’-end cleavage and polyadenylation complex generating mRNA ready for translation. The RNA remaining attached to the elongating RNA polymerase will rapidly be degraded by XRN2 generating no measurable readthrough transcripts and transcription elongation is terminated via the torpedo model (RT not observed). If the transcripts are not spliced, they will not be cut by the 3’-end cleavage complexes, and the RNA polymerases will continue to elongate past the TES site until allosteric termination occur as the polymerases becomes less processive as it passes the TES (RT observed). This “non-productive” transcription is part of genome surveillance to detect and repair DNA damage^80^. **b** Results from this study show that inhibition of splicing increases readthrough transcripts immediately after the TES as do arsenite, hyperosmotic stress and depletion of BRD4 suggesting that 3’-end cleavage is inhibited. Hyperosmotic stress also affects the allosteric termination step as did depletion of MED14 allowing for longer readthrough transcripts. In contrast, depletion in SUPT16H decreases the length of the readthrough transcript suggesting loss of SUPT16H makes allosteric termination more proficient. The figure is based on the scaled readthrough graph in Fig. 2h.

What may be the mechanism behind the increases in the amounts of readthrough transcripts following arsenic and hyperosmotic stress or the depletion of MED14 or BRD4? Our data do not support reduced splicing being responsible since compared to the treatments with the splicing inhibitors, the fraction spliced did not change appreciably after these treatments (Supplementary Fig. 14). Our working model is that the increases observed in readthrough transcripts following arsenic or hyperosmotic stresses or depletion of BRD4 most likely involve a reduced ability to perform 3-end cleavage and polyadenylation (Fig. 7b). Furthermore, hyperosmotic stress and depletion of the Mediator component MED14 suppressed transcription termination of readthrough transcripts while depletion of the FACT complex subunit SUPT16H increased transcription termination (Fig. 7b). More detailed studies are required to uncover the underlying mechanisms by which these cellular stresses and depletion of MED14, BRD4 and SUPT16H regulate readthrough transcripts.

## Materials and Methods

### Cell lines, sequencing, and data mapping

As part of the Deeply Profiled Cell Lines (DPCL) collection, cells were grown and collected in two batches (N = 2 biosamples) for all assays together according to standard methods established for each cell line and then distributed to participating ENCODE consortium labs for completion of experimental protocols and sequencing. Growth conditions of each cell line have been published elsewhere^33^. In addition, experimental guidelines and data standards for most assays can be found on the ENCODE portal (https://www.encodeproject.org/data-standards/).

Sequencing parameters for each sample are available on the ENCODE portal (https://www.encodeproject.org). All data was aligned to GRCh38 and gene-centric analyses used the GENCODE annotation, version 29 (v29, https://www.GENCODEgenes.org/human/release_29.html). The ENCODE Data Coordinating Center (DCC) has developed Uniform Processing Pipelines for most major assay types generated by the consortium. Details about the DCC Uniform Processing Pipelines can be found on the ENCODE portal (https://www.encodeproject.org/pipelines/) and code is available on the ENCODE DCC GitHub (https://github.com/ENCODE-DCC). All cells used throughout this study were routinely cultured at 37℃ in a humidified incubator containing 5% CO2.

### AID-mediated protein depletion and exogenous treatments

HCT116 auxin-inducible degron (AID) cell lines targeting CTCF, RAD21, BRD4, MED14, SUPT16H, SMARCA5 or CDK7 were generated in the laboratory of Dr. Erez Lieberman Aiden at the University of Texas and distributed to participating members of the ENCODE consortium for use in various transcriptional and chromatin assays. Bru-seq was performed using these cell lines as described for the DPCL experiments. Protein depletion was induced by treating the cells with 1 µM 5-Phenyl-1H-indole-3-acetic acid (5-Ph-IAA) (auxin) for 6 hours. Data generated from these experiments are available through the “Protein Knockdown degron” collection on the ENCODE portal (https://www.encodeproject.org).

HCT116-AID cells targeting XRN2, DIS3 and EXOSC10 were a generous gift from Dr.Steven West, University of Exeter, United Kingdom. Cells were cultured in DMEM (GIBCO, 11995065) supplemented with 10% FBS. Auxin (Indole-3-acetic acid, Sigma-Aldrich, I3750-5G-A) was dissolved in 100% ethanol at a stock concentration of 500 mM. For Bru-seq experiments, auxin was added to the culture medium at a final concentration of 500 µM for 1.5 hours before nascent RNA labeling. Bromouridine (Bru, MilliporeSigma, 850187) was then added to a final concentration of 2 mM for 30 minutes, and auxin was present throughout the labeling period, resulting in a total auxin treatment time of 2 hours. The control samples received an equal volume of 100% ethanol. Bru-RNA isolation, library preparation and paired-end sequencing was performed as described previously^39^.

Protein depletion was confirmed by western blotting. Cells were lysed in RIPA buffer containing 50 mM Tris (pH 7.5), 150 mM NaCl, 1% NP-40, 0.1% SDS, 1x Protease Inhibitor cocktail (Roche,11836170001). Samples were prepared in 4x Laemmli sample buffer (Biorad, 1610747) supplemented with β-mercaptoethanol and separated on 4-20% Tris-glycine gradient gels (Bio-rad, 4561094) under denaturing and reducing conditions. Immunoblotting was performed using antibodies against XRN2 (1:1000, Bethyl, A301-103A-M), DIS3 (1:1000, Bethyl, A303-765A-M), and EXOSC10 (1:1000, Abcam, ab50558). Fibrillarin (1:5000, Abcam, ab5821) or β-actin (Sigma-Aldrich, A5441) was used as a loading control.

K562 cells were cultured in suspension in IMDM supplemented with 10% FBS. Meayamycin B (a gift from Dr. Kazunori Koide, University of Pittsburgh) was added to a final concentration of 3 µM. Cells were treated for 6 hours, and nascent RNA was labeled by adding bromouridine to a final concentration of 2 mM during the last 30 minutes of treatment. Cells were then harvested for Bru-seq or subjected to a chase for BruChase-seq. For the chase, cells were collected by centrifugation, washed once with PBS, and resuspended in an equal volume of conditioned media containing Meayamycin B. Uridine (MilliporeSigma, U3750) was added to a final concentration of 20 mM, and the cells were incubated for an additional 2 or 6 hours. Control cells were processed similarly, except that DMSO was added to the cells instead of Meayamycin B. Bru-RNA isolation, library preparation and single-end sequencing was performed as previously described^37^.

For the remaining experimental conditions, hTERT immortalized human diploid foreskin fibroblasts (HF1) (a gift from Dr. Mary Davis, Department of Radiation Oncology, University of Michigan, Ann Arbor) were grown in Minimal Essential Medium with 10% FBS^36,37^, and treated with 100 nM pladienolide-B (Plad-B, EMD Millipore, 530196) for 1 hour. HOS cells were cultured in DMEM supplemented with 10% FBS. Cells were dissociated using 3 ml of 0.25% trypsin for 5 minutes and seeded at a density of 1 million cells per 10 cm dish. Cells were then treated with 10 µM Vorinostat (Selleckchem, S1047) for 16 hours. U2OS cells were maintained in McCoy’s 5a medium with 10% FBS as described previously^86^. For the heat shock treatment, U2OS cells were incubated at 42℃ and harvested after an hour. To induce oxidative stress, sodium arsenite was added to U2OS cells at a final concentration of 0.5 mM for 1 hour. For the hyperosmotic condition, U2OS cells were treated with 300 mM sodium chloride for 1 hour. Nascent RNA was labeled with 2 mM Bru during the final 30 minutes of each treatment. Bru-RNA isolation, library preparation and paired-end sequencing was performed as described previously^39^.

### Genome coverage calculations

To calculate the proportion of the genome covered by at least one uniquely mapping sequencing read for each assay and cell line, we generated unnormalized, stranded bigwigs from alignments using deeptools^87^ bamCoverage for each sample using the following parameters: *--normalizeUsing None --minMappingQuality 255 --filterRNAstrand forward/reverse --binSize 1 --skipNAs*. We then merged bigwigs per strand using UCSC utilities bigWigMerge^88^ for each application that assessed the coverage of more than one sample. The resulting bedgraph files were then sorted (UCSC utilities bedSort) and reverted to bigwigs (UCSC utilities bedGraphToBigWig). Coverages were calculated for each of the canonical chromosomes (1-22, X, Y) per strand from merged bigwigs using the pyBigWig (deeptools) bw.stats(type=“coverage”), and the total bases covered on both strands was calculated for all chromosomes to yield a signal fraction of bases covered per bigwig.

For genic, intergenic, and exonic coverage calculations, merged bigwigs were generated as described above for Bru-seq, BruUV-seq, and total RNA-seq using only the 24 libraries for which we had data in all three assays (8 samples are missing from DPCL total RNA-seq), to keep these data as consistent as possible. Similarly, these data are only shown for the 8 cell lines (16 libraries) which had two biological replicates in all three assays. To calculate the fractions of bases covered in the genic versus intergenic regions of the genome, we first obtained an annotation of these regions (.bed format). We merged (bedtools merge^89^) gene boundaries from the GENCODE v29 basic annotation in a stranded fashion to establish the genic region, and then assigned the remaining regions, up to the length of each chromosome (GRCh38), as intergenic. To obtain the exonic regions, we first created a collapse exon annotation (GENCODE v29 basic) per gene using a custom python script (available on GitHub) and then merged any overlapping exonic regions on the same strand with bedtools as described above for genes. The coverages in each of these regions were calculated with pyBigWig.stats(type=“coverage”) as described above and the total bases covered was calculated per region, yielding 3 coverage values per bigwig (genic, intergenic, exonic).

Finally, genic versus intergenic counts were calculated for each bigwig using featureCounts (subread)^90^ using the following commands: *-F SAF -p (excluded for total RNA-seq) -Q 255 -s 2 --countReadPairs (excluded for total RNA-seq)*.

Counts were then summed per region, and the fraction of total counts was calculated using the total number of uniquely mapping reads per bigwig (sum of all region counts). To compare cell lines, Pearson’s correlations (r) were calculated from a counts matrix and hierarchical clustering was performed using *pheatmap(clustering_distance_rows = “correlation”, clustering_distance_cols = “correlation”, clustering_method = “complete”, cluster_rows = T, cluster_cols = T)*.

### RT segment (RTseg) identification

To infer regions of readthrough transcription, we performed genome segmentation using a previously described Hidden Markov model (HMM) with 10 logarithmically distributed output states. Here we used 250bp binned RPKM values as an input to this model, which establishes the inferred transcription states of genome segments (HMM index 0-9), and excluded segments with an HMM index less than three. We then merged adjacent segments with equal or decreasing HMM indices and intersected these merged segments with gene transcription end sites (TESs). Using the GENCODE v29 annotation, one TES was obtained per gene by selecting the 3’-most end site from all annotated gene isoforms. Each segment that overlapped multiple TESs was assigned to the most upstream TES that it overlapped, and the 5’-coordinate of all segments was defined by the TES of the associated gene. These segments were considered possible readthrough segments for their associated gene and were subject to further filtering. Readthrough segments were not reported for genes that were less than 1kb in length or were lowly expressed (RPKM < 0.25). These length and expression cutoffs were taken due to the inability to accurately resolve Bru-seq signal and RT segments at small and lowly expressed genes. Coverage over readthrough segments was obtained, and all segments with fewer than 10 counts were not reported. The remaining segments were reported as readthrough segments (RTsegs) and were subject to downstream classification and refinement based on their overlaps with annotated genes. RTsegs were identified for each sample individually but were merged to create a cell line specific annotation of RT regions for most analyses. To merge replicates, the longest segment from either sample was retained per gene, relying on the assumption that each replicate would have identical RT potential downstream of a particular gene.

### RT versus NRT gene definitions

Due to the restriction of our analysis to genes that are greater than 1 kb in length we assayed a total of 38006 genes (GENCODE v29) for read through. RT genes were identified as genes that were assigned an RTseg in at least one cell line (18244) and NRT genes were not assigned an RTseg in any cell line (19762). These analyses are independent of the length of a RTseg. However, to remove the possibility that technical ambiguity could confound our results, we also obtained a subset of stringent NRT genes (N = 239). Stringent NRT genes: 1.) were required to be expressed (RPKM > 0.25) in at least one cell line, 2.) have little to no RNA signal in the 5 kb downstream of their TES (<10 Bru-seq reads), 3.) do not have another gene within 5 kb downstream, and 4.) their TESs are not overlapped by an RTseg coming from an upstream gene on the same strand. Gene expression was not enforced in all cell lines due to the presence of very few ubiquitously expressed genes that met all other listed requirements. Stringent RT genes are a subset of 239 genes where: 1.) at least 1 cell line does not overlap a downstream gene on the same strand, 2.) no cell lines have ambiguous RTseg classifications, and 3.) do not have another gene downstream within 5kb. The distribution of biotypes between the two subsets–and the overall distribution of RT gene biotypes–was kept as similar as possible to remove the potential for differences between coding or non-coding gene features to influence our results in either subset. The distribution of gene lengths between the two stringent subsets were also confirmed to be similar. In addition to the stringent subset, all results were verified using a subset of only protein-coding genes due to the imbalance of biotypes in the full RT/NRT gene sets and to more closely resemble the results of previous studies which have typically attributed RT transcripts to protein-coding genes exclusively. These gene lists were used for all analyses comparing nascent RT transcription to known steady-state homeostatic or stress-induced RT transcripts.

### Scaled RT signal calculation

Nascent readthrough signal was evaluated from scaled binned counts (250 bp bins) downstream of an RT gene TES. These binned counts were either obtained for the entire RTseg identified in our segmentation analysis, or for a fixed 17377 bp region downstream of RT genes. RT signal length was evaluated using binned counts for the full-length segments, and comparisons between RT signal patterns found across cell lines were performed using the fixed 17.3 kb regions. For all RT genes, the binned counts in the RT region were scaled to the signal at the end of the gene to allow comparison of RT signal patterns between genes and cell lines. To do this, we obtained the counts of four 250 bp bins upstream of the TES (∼1 kb into the gene body) and divided the binned counts in the readthrough region by the maximum binned count value at the end of the gene. Because multi-isoform genes, particularly those with multiple annotated TESs, could result in inaccurate measurements of the end-of-gene signal used for scaling, we restricted this analysis to single-isoform genes and genes where all annotated TESs are within 250 bp (1 bin size) of each other. These scaled RT signals were then further filtered for downstream analyses as needed.

### RT signal scaling for exogenous stresses and knockdown conditions

To determine signal profiles downstream of the Transcription End Site (TES) for various conditions, we first generated strand-specific bigWig files from alignment files using deeptools (v3.5.5) with the following options: *bamCoverage -b --binSize 1 --normalizeUsing RPKM -- exactScaling --minMappingQuality 255 --filterRNAstrand forward* or *reverse*. Protein-coding genes ≥ 10 kb were selected, excluding genes with any annotated feature (any biotype, GRCh38, GENCODE v29) within 40 kb downstream of the TES on either strand (number of genes = 1462). For each stranded bigwig, coverage over the genomic regions of interest was quantified separately for plus- and minus-strand genes using deepTools *computeMatrix reference-point -S --referencePoint center --upstream 10000 --downstream 40000 --binSize 50 --sortRegions descend --sortUsing median --averageTypeBins median*. For each sample, sense and antisense signals were computed per strand, and binned scores were generated with deepTools *plotProfile -m --averageType mean*. The scores for each 50 bp bin in the downstream 40 kb region were scaled to the 1 kb region upstream of the TES to generate scaled bin values. This normalization was performed only for the sense strand signal. The resulting scaled binned values were used to produce the final metagene plots in R (R/ggplot2).

### Approximate distances traveled by RT signal

Scaled counts from full-length RTsegs were used to calculate approximate distances or lengths to a particular signal level (number of bins * bin length) for most RT genes. RT genes were excluded from these analyses if their RTseg length was less than 250 bp or if they had RTsegs with ambiguous classifications. The number of bins to each signal level was determined by setting a scaled signal threshold (e.g. 0, 0.01, 0.1, 0.5) and counting how many bins it took until the signal was maintained at or below that threshold for four consecutive bins (1 kb). The distribution of approximate RT signal lengths to zero scaled counts was reported and compared with RTseg lengths and the approximate distance to different signal levels (50, 10, 1, 0% of the gene signal) allowed quantification of the different signal trajectories downstream of RT genes (Fig. 2I). A threshold of zero counts was considered appropriate for these analyses based on the use of segmentation data to define the assayed regions before counting, thus we do not expect to be capturing noise.

### Comparisons of scaled RT signals between cell lines

To compare RT signals for the same gene across different cell lines, we obtained scaled counts as described above for 17377 bp regions (equal to the median length of all RT segments). In addition to the filters applied to all RT signal regions, we removed RT genes that: 1.) had RTseg lengths less than the fixed region (17.3 kb), 2.) had scaled RT signals where one or more bins had a higher count value than the gene body maximum (i.e. scaled signal must be between 0 and 1), and 3.) that had a downstream gene on the same strand within 17.3 kb. These measures were taken, to avoid ambiguities in the downstream RT signal that are attributable to gene expression that is unrelated to the parent gene, however it is still possible that signal artifacts could exist in this subset (e.g. highly expressed enhancers or unannotated genes). This final subset of fixed-length RT signals was then aggregated per cell line, and each RT signal was normalized to the median aggregated signal by subtracting the median value per cell line from each bin. The normalized signals from this analysis were then used to describe the variance between RT signals, where positive values indicate binned scaled counts higher than the median and vice versa.

### Assessing the chromatin landscape of RT/NRT genes

To compare the chromatin landscape at the ends and downstream of nascent RT genes, we utilized several ENCODE4 datasets describing histone modifications, CTCF binding, chromatin accessibility, and units of 3D chromatin organization. Peaks of ENCODE4 ChIP-seq signal were obtained for the chromatin binding protein CTCF and the following histone modifications: H3K27me3, H3K27ac, H3K4me1, H3K4me3, H3K36me3, and H3K79me2 (.bed). In addition, we used peaks called from DPCL ATAC-seq and DNase-seq data, and loop anchors from DPCL POLR2A ChIA-PET and intact Hi-C loops (.bedpe). Finally, chromatin domain boundaries (TAD boundaries) and A/B compartments (.bed) were obtained from DPCL intact Hi-C data. All these data describing various features of the chromatin landscape will be referred to as our chromatin regions-of-interest when discussed together throughout the rest of this section.

For each cell line for which we had each chromatin data type, we counted the frequency that the 1 kb upstream (TES-1kb) and 5 kb region downstream (TES+5kb) of each gene TES was overlapped by a chromatin region-of-interest using bedtools (intersect for .bed files and pairToBed for .bedpe files). As with all RT analyses, genes were required to be greater than 1 kb in length. Genes were grouped based on the presence (ROI) or absence (noROI) of an overlapping chromatin region-of-interest, either in each cell line individually or in any cell line. This left us with a binary grouping describing the patterns of chromatin region-of-interest on a per cell line basis or for all cell lines combined. As before, we split our overall gene list into RT and NRT genes but removed any gene that had an annotated gene on either strand (sense or antisense) within 7 kb of its TES. This resulted in total RT and NRT subsets of 6187 and 8313, respectively. For cell line specific RT/NRT subsets, we obtained all RT genes found in a given cell line and generated a random sample of NRT genes of an equal size using the sample() function in base R.

We then performed a Fisher’s exact test to compare our binary groups (RT/NRT and ROI/noROI) for each cell line individually and all cell lines combined. P-values were corrected for multiple-comparisons using the Bonferroni test, and adjusted p-values were used to report significant differences between groups. Bar graphs were used to summarize the fraction of all RT and NRT genes that had an overlapping chromatin region-of-interest in any cell line. To summarize cell line specific trends between RT/NRT genes and chromatin regions-of-interest, we performed a tetrachoric correlation96 (https://john-uebersax.com/stat/tetra.htm) using the tetrachoric() function from the psych package in R. between our binary groups (RT/NRT and ROI/noROI) and reported the correlation coefficient (rtet) in a heatmap along with Fisher’s exact adjusted p-values as calculated above. This analysis was performed for all RT/NRT genes.

### Sequence analysis of RT/NRT genes

To understand if there are sequence motifs that demarcate RT genes, such as higher GC content, we obtained the sequences (GRCh38) in the 1 kb upstream (TES-1kb) and 5 kb downstream (TES+5kb) of gene TESs using bedtools getfasta (-s -bedOut). For sequence analyses, we required that genes had no annotated gene on either strand within 7 kb downstream of its TES. Using custom python scripts to parse sequence strings, we quantified the occurrences of each nucleotide (A, T, C, G) and calculated the GC-content of each region flanking the TES: (G+C)/(G+C+A+T). The distribution of GC-content was then reported for RT versus NRT genes both upstream and downstream of the TES (TES-1kb, TES+5kb).

### Calculation of the proportion of spliced reads

A single-end (SE) read, or paired-end (PE) read pair was classified as spliced if the CIGAR string of the SE read or either mate of the PE read pair contained an “N”. The fraction of spliced SE reads or PE read pairs was calculated relative to all uniquely mapped SE reads or PE read pairs (MAPQ = 255) on the standard chromosomes.

## Supporting information

Supplemental Figgures 1-14

## Data Availability

The Bru-seq datasets used in this study are available through the ENCODE portal (https://www.encodeproject.org/): Data for the 16 cell lines can be found in the “Deeply Profiled Cell Lines” collection, whereas the HCT116-AID datasets targeting various chromatin modifiers are included in the “Protein Knockdown Degron” collection. The U2OS dataset is available through GEO under accession number GSE319125, while all other datasets are available under accession number GSE352755. Custom analysis scripts developed for this manuscript are available on GitHub at https://github.com/LjungmanLab/ENCODE-DPCL-readthrough.

## Acknowledgements

We thank the personnel at University of Michigan Advanced Genomics Core for professional technical assistance with sequencing. We thank Nivea Vydiswaran for technical assistance with the cell stress experiments. We also thank Drs. JoAnn Sekiguchi and Steve Parker and members of the ENCODE Consortium for valuable input into this study.

## Author contributions

A.McS. performed and analyzed Bru-seq data displayed in Figures 1, 2 and supplementary Fig. 1 as well as Supplementary Dataset 1 and 2 and contributed to the writing of the manuscript. I.V.N. performed experiments displayed in Fig. 5, Supplementary Fig. 7-10 and Supplementary Data sets 9-11. She also calculated scaled readthrough aggregate plots for all samples and contributed to the writing of the manuscript. M.P. grew the 16 cell lines part of “Deeply Profiled Cell Lines” collection for experiments shown in Figs 1 and 2 and performed all the Bru-seq experiments including capturing of the Bru-labeled RNA and library preparations for deep sequencing. H.B. contributed to cell culturing and data presented in Supplementary Fig. 6 and Supplementary Dataset 8, M.A., B.M., K.B., and N.T.Y. contributed to computational analyses throughout the study, J.P. performed the experiments shown in Fig. 4 and Supplementary Dataset 7, J.D. and S.P. performed the experiments displayed in Fig. 3 and Supplementary Figs. 2-4, and Supplementary Datasets 3-5, E.S. and P.G. performed the data presented in Supplementary Fig. 5 and Supplementary Data set 6, E.A.L. performed the Auxin experiments shown in Fig. 6 and Supplementary Figs. 8, 11-13 and Supplementary Datasets 12-18. T.E.W. designed the Bru-seq analysis pipeline used for the analysis of all the Bru-seq experiments, M.L. invented the Bru-seq techniques, performed analyses, made figures and wrote the manuscript. All authors discussed and edited the manuscript.

## References

1. Girbig, M., Misiaszek, A.D. & Muller, C.W. Structural insights into nuclear transcription by eukaryotic DNA-dependent RNA polymerases. Nat Rev Mol Cell Biol 23, 603–622 (2022).

2. Nojima, T. & Proudfoot, N.J. Mechanisms of lncRNA biogenesis as revealed by nascent transcriptomics. Nat Rev Mol Cell Biol 23, 389–406 (2022).

3. Jonkers, I. & Lis, J.T. Getting up to speed with transcription elongation by RNA polymerase II. Nat Rev Mol Cell Biol 16, 167–77 (2015).

4. Chen, F.X., Smith, E.R. & Shilatifard, A. Born to run: control of transcription elongation by RNA polymerase II. Nat Rev Mol Cell Biol 19, 464–478 (2018).

5. Porrua, O., Boudvillain, M. & Libri, D. Transcription Termination: Variations on Common Themes. Trends Genet 32, 508–522 (2016).

6. Kuehner, J.N., Pearson, E.L. & Moore, C. Unravelling the means to an end: RNA polymerase II transcription termination. Nat Rev Mol Cell Biol 12, 283–94 (2011).

7. Richard, P. & Manley, J.L. Transcription termination by nuclear RNA polymerases. Genes Dev 23, 1247–69 (2009).

8. Elkon, R., Ugalde, A.P. & Agami, R. Alternative cleavage and polyadenylation: extent, regulation and function. Nat Rev Genet 14, 496–506 (2013).

9. Neve, J., Patel, R., Wang, Z., Louey, A. & Furger, A.M. Cleavage and polyadenylation: Ending the message expands gene regulation. RNA Biol 14, 865–890 (2017).

10. Eaton, J.D., Francis, L., Davidson, L. & West, S. A unified allosteric/torpedo mechanism for transcriptional termination on human protein-coding genes. Genes Dev 34, 132–145 (2020).

11. Cortazar, M.A. et al. Xrn2 substrate mapping identifies torpedo loading sites and extensive premature termination of RNA pol II transcription. Genes Dev 36, 1062–1078 (2022).

12. Rodriguez-Molina, J.B., West, S. & Passmore, L.A. Knowing when to stop: Transcription termination on protein-coding genes by eukaryotic RNAPII. Mol Cell 83, 404–415 (2023).

13. Sousa-Luis, R. et al. POINT technology illuminates the processing of polymerase-associated intact nascent transcripts. Mol Cell 81, 1935–1950 e6 (2021).

14. Mo, W. et al. Landscape of transcription termination in Arabidopsis revealed by single-molecule nascent RNA sequencing. Genome Biol 22, 322 (2021).

15. Alfonso-Gonzalez, C. & Hilgers, V. Elucidating the coordination of RNA processing using short-read and long-read RNA-sequencing methods. Nat Rev Mol Cell Biol 27, 194–212 (2026).

16. Skourti-Stathaki, K., Kamieniarz-Gdula, K. & Proudfoot, N.J. R-loops induce repressive chromatin marks over mammalian gene terminators. Nature 516, 436–9 (2014).

17. Grzechnik, P., Tan-Wong, S.M. & Proudfoot, N.J. Terminate and make a loop: regulation of transcriptional directionality. Trends Biochem Sci 39, 319–27 (2014).

18. Fong, N. et al. Effects of Transcription Elongation Rate and Xrn2 Exonuclease Activity on RNA Polymerase II Termination Suggest Widespread Kinetic Competition. Mol Cell 60, 256–67 (2015).

19. Muniz, L. et al. Control of Gene Expression in Senescence through Transcriptional Read-Through of Convergent Protein-Coding Genes. Cell Rep 21, 2433–2446 (2017).

20. Caldas, P. et al. Transcription readthrough is prevalent in healthy human tissues and associated with inherent genomic features. Commun Biol 7, 100 (2024).

21. Grosso, A.R. et al. Pervasive transcription read-through promotes aberrant expression of oncogenes and RNA chimeras in renal carcinoma. Elife 4(2015).

22. Rutkowski, A.J. et al. Widespread disruption of host transcription termination in HSV-1 infection. Nat Commun 6, 7126 (2015).

23. Vilborg, A., Passarelli, M.C., Yario, T.A., Tycowski, K.T. & Steitz, J.A. Widespread Inducible Transcription Downstream of Human Genes. Mol Cell 59, 449–61 (2015).

24. Vilborg, A. et al. Comparative analysis reveals genomic features of stress-induced transcriptional readthrough. Proc Natl Acad Sci U S A 114, E8362–E8371 (2017).

25. Rosa-Mercado, N.A. et al. Hyperosmotic stress alters the RNA polymerase II interactome and induces readthrough transcription despite widespread transcriptional repression. Mol Cell 81, 502–513 e4 (2021).

26. Jalihal, A.P. et al. Multivalent Proteins Rapidly and Reversibly Phase-Separate upon Osmotic Cell Volume Change. Mol Cell 79, 978–990 e5 (2020).

27. Cardiello, J.F., Goodrich, J.A. & Kugel, J.F. Heat Shock Causes a Reversible Increase in RNA Polymerase II Occupancy Downstream of mRNA Genes, Consistent with a Global Loss in Transcriptional Termination. Mol Cell Biol 38(2018).

28. Abrisch, R.G., Eidem, T.M., Yakovchuk, P., Kugel, J.F. & Goodrich, J.A. Infection by Herpes Simplex Virus 1 Causes Near-Complete Loss of RNA Polymerase II Occupancy on the Host Cell Genome. J Virol 90, 2503–13 (2015).

29. Bauer, D.L.V. et al. Influenza Virus Mounts a Two-Pronged Attack on Host RNA Polymerase II Transcription. Cell Rep 23, 2119–2129 e3 (2018).

30. Hennig, T. et al. HSV-1-induced disruption of transcription termination resembles a cellular stress response but selectively increases chromatin accessibility downstream of genes. PLoS Pathog 14, e1006954 (2018).

31. Walsh, K.E., Goodrich, J.A. & Kugel, J.F. CPSF73 activation and 3’ RNA polymerase II pausing are lost during readthrough transcription after heat shock. Cell Rep 45, 117039 (2026).

32. Agostini, F., Zagalak, J., Attig, J., Ule, J. & Luscombe, N.M. Intergenic RNA mainly derives from nascent transcripts of known genes. Genome Biol 22, 136 (2021).

33. Narayanan, I.V. et al. Isoform- and pathway-specific regulation of post-transcriptional RNA processing in human cells. Genome Res (2026).

34. Rosa-Mercado, N.A. & Steitz, J.A. Who let the DoGs out? - biogenesis of stress-induced readthrough transcripts. Trends Biochem Sci 47, 206–217 (2022).

35. Bedi, K., Paulsen, M.T., Wilson, T.E. & Ljungman, M. Characterization of novel primary miRNA transcription units in human cells using Bru-seq nascent RNA sequencing. NAR Genom Bioinform 2, lqz014 (2020).

36. Paulsen, M.T. et al. Use of Bru-Seq and BruChase-Seq for genome-wide assessment of the synthesis and stability of RNA. Methods 67, 45–54 (2014).

37. Paulsen, M.T. et al. Coordinated regulation of synthesis and stability of RNA during the acute TNF-induced proinflammatory response. Proc Natl Acad Sci U S A 110, 2240–5 (2013).

38. Schwalb, B. et al. TT-seq maps the human transient transcriptome. Science 352, 1225–8 (2016).

39. Magnuson, B. et al. CDK12 regulates co-transcriptional splicing and RNA turnover in human cells. iScience 25, 105030 (2022).

40. Morishita, K., Watanabe, K., Naguro, I. & Ichijo, H. Sodium ion influx regulates liquidity of biomolecular condensates in hyperosmotic stress response. Cell Rep 42, 112315 (2023).

41. Ruiz-Ramos, R., Lopez-Carrillo, L., Rios-Perez, A.D., De Vizcaya-Ruiz, A. & Cebrian, M.E. Sodium arsenite induces ROS generation, DNA oxidative damage, HO-1 and c-Myc proteins, NF-kappaB activation and cell proliferation in human breast cancer MCF-7 cells. Mutat Res 674, 109–15 (2009).

42. Rainey, N.E., Armand, A.S. & Petit, P.X. Sodium arsenite and arsenic trioxide differently affect the oxidative stress of lymphoblastoid cells: An intricate crosstalk between mitochondria, autophagy and cell death. PLoS One 19, e0302701 (2024).

43. Grant, S., Easley, C. & Kirkpatrick, P. Vorinostat. Nat Rev Drug Discov 6, 21–2 (2007).

44. Lee, J.H., Choy, M.L., Ngo, L., Foster, S.S. & Marks, P.A. Histone deacetylase inhibitor induces DNA damage, which normal but not transformed cells can repair. Proc Natl Acad Sci U S A 107, 14639–44 (2010).

45. Herzel, L., Straube, K. & Neugebauer, K.M. Long-read sequencing of nascent RNA reveals coupling among RNA processing events. Genome Res 28, 1008–1019 (2018).

46. Drexler, H.L., Choquet, K. & Churchman, L.S. Splicing Kinetics and Coordination Revealed by Direct Nascent RNA Sequencing through Nanopores. Mol Cell 77, 985–998 e8 (2020).

47. Fair, B. et al. Global impact of unproductive splicing on human gene expression. Nat Genet 56, 1851–1861 (2024).

48. Bedi, K. et al. Co-transcriptional splicing efficiencies differ within genes and between cell types. RNA (2021).

49. Tellier, M., Maudlin, I. & Murphy, S. Transcription and splicing: A two-way street. Wiley Interdiscip Rev RNA 11, e1593 (2020).

50. Custodio, N. et al. Inefficient processing impairs release of RNA from the site of transcription. EMBO J 18, 2855–66 (1999).

51. Kaida, D. et al. Spliceostatin A targets SF3b and inhibits both splicing and nuclear retention of pre-mRNA. Nat Chem Biol 3, 576–83 (2007).

52. Kotake, Y. et al. Splicing factor SF3b as a target of the antitumor natural product pladienolide. Nat Chem Biol 3, 570–5 (2007).

53. West, S., Gromak, N. & Proudfoot, N.J. Human 5’ --> 3’ exonuclease Xrn2 promotes transcription termination at co-transcriptional cleavage sites. Nature 432, 522–5 (2004).

54. Davidson, L. et al. Rapid Depletion of DIS3, EXOSC10, or XRN2 Reveals the Immediate Impact of Exoribonucleolysis on Nuclear RNA Metabolism and Transcriptional Control. Cell Rep 26, 2779–2791 e5 (2019).

55. Lemay, J.F. et al. The RNA exosome promotes transcription termination of backtracked RNA polymerase II. Nat Struct Mol Biol 21, 919–26 (2014).

56. Lemay, J.F. & Bachand, F. Fail-safe transcription termination: Because one is never enough. RNA Biol 12, 927–32 (2015).

57. Rao, S.S. et al. A 3D Map of the Human Genome at Kilobase Resolution Reveals Principles of Chromatin Looping. Cell 159, 1665–80 (2014).

58. Phillips, J.E. & Corces, V.G. CTCF: master weaver of the genome. Cell 137, 1194–211 (2009).

59. Fudenberg, G. et al. Formation of Chromosomal Domains by Loop Extrusion. Cell Rep 15, 2038–49 (2016).

60. Ganji, M. et al. Real-time imaging of DNA loop extrusion by condensin. Science 360, 102–105 (2018).

61. Cheng, H., Zhang, N. & Pati, D. Cohesin subunit RAD21: From biology to disease. Gene 758, 144966 (2020).

62. Barisic, D., Stadler, M.B., Iurlaro, M. & Schubeler, D. Mammalian ISWI and SWI/SNF selectively mediate binding of distinct transcription factors. Nature 569, 136–140 (2019).

63. Bomber, M.L. et al. Human SMARCA5 is continuously required to maintain nucleosome spacing. Mol Cell 83, 507–522 e6 (2023).

64. Belew, M.D., Chen, J. & Cheng, Z. Emerging roles of cyclin-dependent kinase 7 in health and diseases. Trends Mol Med 31, 138–151 (2025).

65. Putlyaev, E.V., Ibragimov, A.N., Lebedeva, L.A., Georgiev, P.G. & Shidlovskii, Y.V. Structure and Functions of the Mediator Complex. Biochemistry (Mosc) 83, 423–436 (2018).

66. Lu, C. et al. Mediator regulates transcriptional termination through crosstalk with pre-mRNA 3’ end processing factors. Mol Cell 85, 2147–2164 e10 (2025).

67. Dhalluin, C. et al. Structure and ligand of a histone acetyltransferase bromodomain. Nature 399, 491–6 (1999).

68. Jang, M.K. et al. The bromodomain protein Brd4 is a positive regulatory component of P-TEFb and stimulates RNA polymerase II-dependent transcription. Mol Cell 19, 523–34 (2005).

69. Arnold, M., Bressin, A., Jasnovidova, O., Meierhofer, D. & Mayer, A. A BRD4-mediated elongation control point primes transcribing RNA polymerase II for 3’-processing and termination. Mol Cell 81, 3589–3603 e13 (2021).

70. Orphanides, G., LeRoy, G., Chang, C.H., Luse, D.S. & Reinberg, D. FACT, a factor that facilitates transcript elongation through nucleosomes. Cell 92, 105–16 (1998).

71. Orphanides, G., Wu, W.H., Lane, W.S., Hampsey, M. & Reinberg, D. The chromatin-specific transcription elongation factor FACT comprises human SPT16 and SSRP1 proteins. Nature 400, 284–8 (1999).

72. Jamai, A., Puglisi, A. & Strubin, M. Histone chaperone spt16 promotes redeposition of the original h3-h4 histones evicted by elongating RNA polymerase. Mol Cell 35, 377–83 (2009).

73. Geisberg, J.V., Moqtaderi, Z. & Struhl, K. Chromatin regulates alternative polyadenylation via the RNA polymerase II elongation rate. Proc Natl Acad Sci U S A 121, e2405827121 (2024).

74. Djebali, S. et al. Landscape of transcription in human cells. Nature 489, 101–8 (2012).

75. Windhager, L. et al. Ultrashort and progressive 4sU-tagging reveals key characteristics of RNA processing at nucleotide resolution. Genome Res (2012).

76. Singh, J. & Padgett, R.A. Rates of in situ transcription and splicing in large human genes. Nat Struct Mol Biol 16, 1128–33 (2009).

77. Herzel, L., Ottoz, D.S.M., Alpert, T. & Neugebauer, K.M. Splicing and transcription touch base: co-transcriptional spliceosome assembly and function. Nat Rev Mol Cell Biol 18, 637–650 (2017).

78. Tilgner, H. et al. Deep sequencing of subcellular RNA fractions shows splicing to be predominantly co-transcriptional in the human genome but inefficient for lncRNAs. Genome Res 22, 1616–25 (2012).

79. Rogalska, M.E. et al. Transcriptome-wide splicing network reveals specialized regulatory functions of the core spliceosome. Science 386, 551–560 (2024).

80. Ljungman, M. Transcription and genome integrity. DNA Repair (Amst) 118, 103373 (2022).

81. Shenasa, H. & Bentley, D.L. Pre-mRNA splicing and its cotranscriptional connections. Trends Genet 39, 672–685 (2023).

82. Davidson, L. & West, S. Splicing-coupled 3’ end formation requires a terminal splice acceptor site, but not intron excision. Nucleic Acids Res 41, 7101–14 (2013).

83. Dye, M.J. & Proudfoot, N.J. Terminal exon definition occurs cotranscriptionally and promotes termination of RNA polymerase II. Mol Cell 3, 371–8 (1999).

84. Yesbolatova, A. et al. The auxin-inducible degron 2 technology provides sharp degradation control in yeast, mammalian cells, and mice. Nat Commun 11, 5701 (2020).

85. Natsume, T., Kiyomitsu, T., Saga, Y. & Kanemaki, M.T. Rapid Protein Depletion in Human Cells by Auxin-Inducible Degron Tagging with Short Homology Donors. Cell Rep 15, 210–218 (2016).

86. Dudley, J.N., et al. Divergent condensates tune transcriptional responses during stress. bioRxiv (2026).

87. Ramirez, F. et al. deepTools2: a next generation web server for deep-sequencing data analysis. Nucleic Acids Res 44, W160–5 (2016).

88. Kent, W.J., Zweig, A.S., Barber, G., Hinrichs, A.S. & Karolchik, D. BigWig and BigBed: enabling browsing of large distributed datasets. Bioinformatics 26, 2204–7 (2010).

89. Quinlan, A.R. & Hall, I.M. BEDTools: a flexible suite of utilities for comparing genomic features. Bioinformatics 26, 841–2 (2010).

90. Liao, Y., Smyth, G.K. & Shi, W. featureCounts: an efficient general purpose program for assigning sequence reads to genomic features. Bioinformatics 30, 923–30 (2014).

