## Supplemental Figgures 1-14 for "Characterizing nascent readthrough transcripts in human cells"

### Supplementary Figures

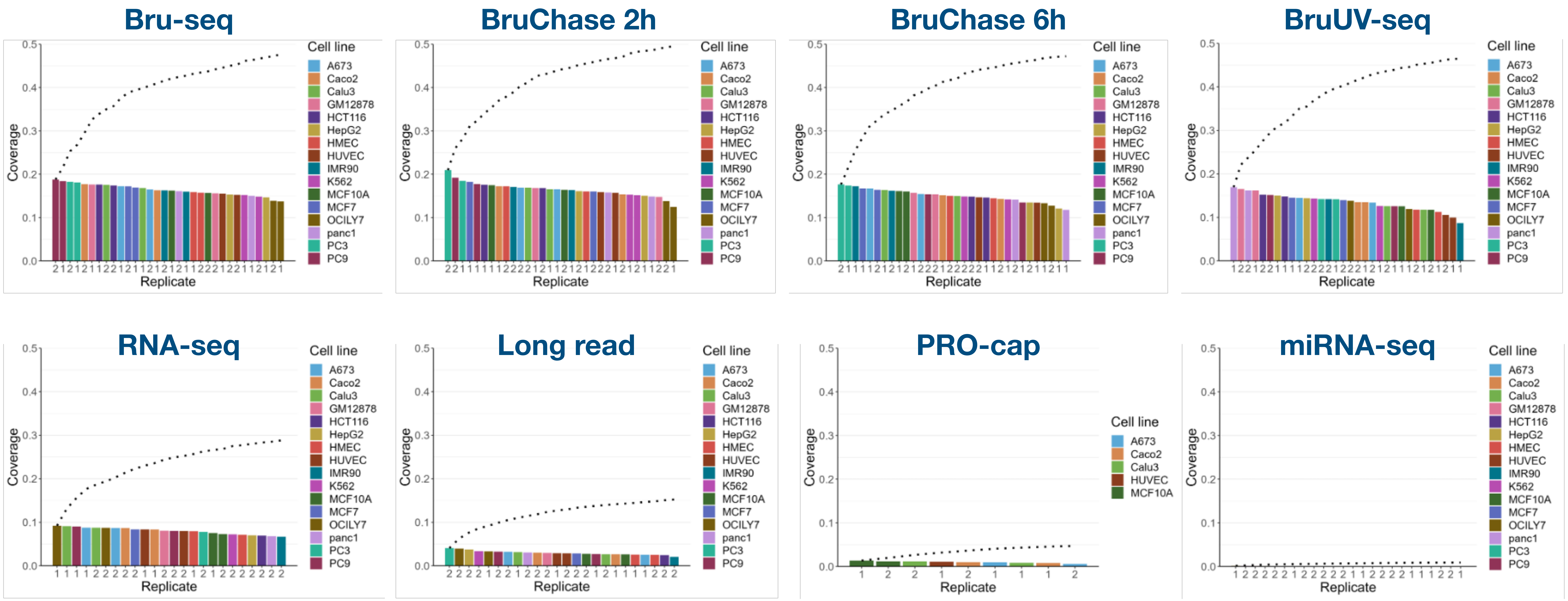


**Supplementary Fig. 1.** Cumulative fraction of bases covered. The cumulative fraction of bases covered by all cell lines for each sequencing method. Each dot representing the cumulative fraction after the fraction of each sample was added.

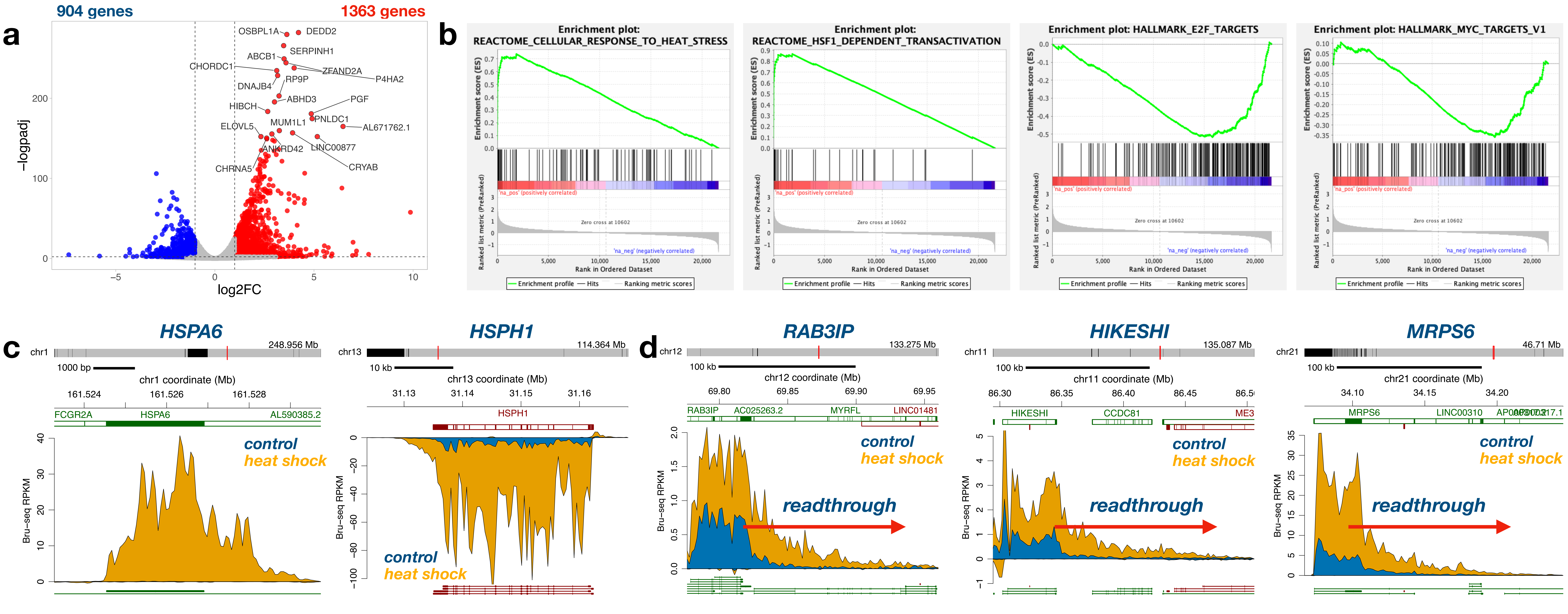


**Supplementary Fig. 2.** Effects of heat shock on gene transcription and readthrough transcripts. **a** Volcano plot comparing nascent transcription of genes of heat shock and mock-treated U2OS cells. Red color indicates induction, blue indicates repression. **b** Gene set enrichment analysis (GSEA) showing two top upregulated and two top downregulated gene sets. **c** Validation of effective heat shock treatment with induction of HSPA6 and HSPH1 genes. Reads from mock-treated cells in blue and heat shock treated cells in yellow. **d** Examples of three genes showing heat shock-induced readthrough transcripts (yellow).

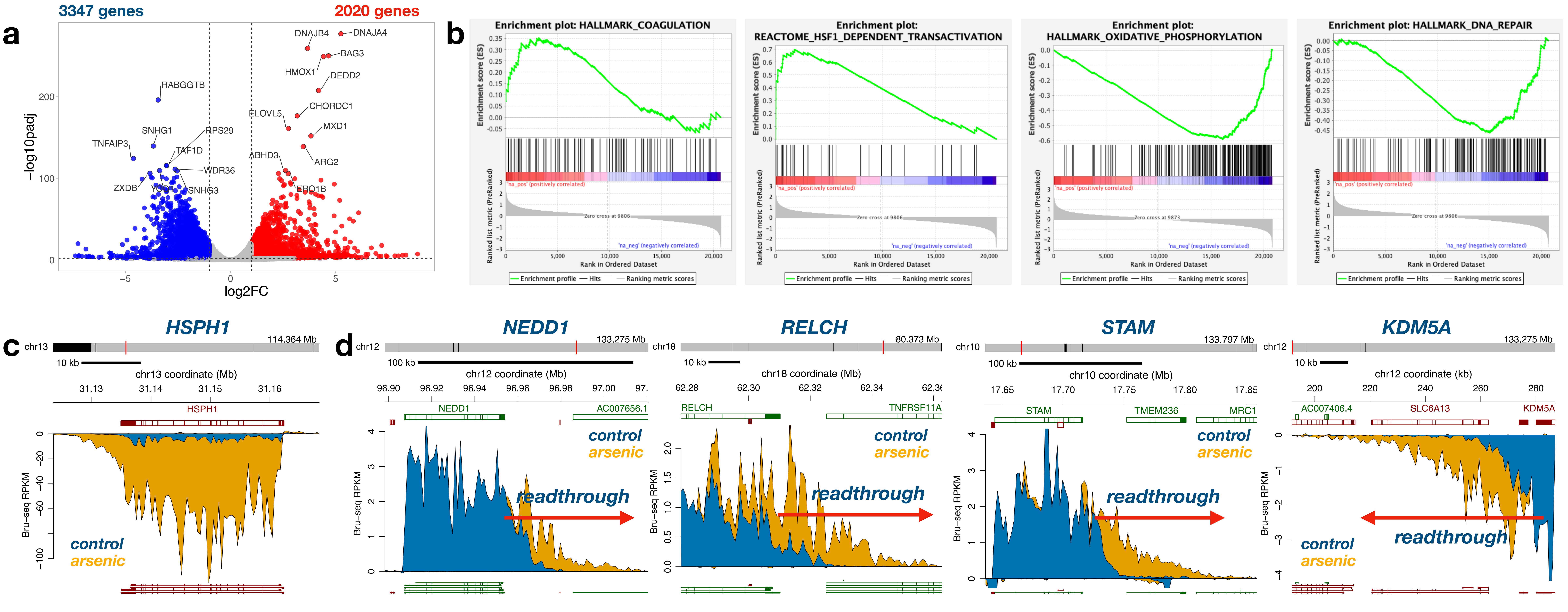


**Supplementary Fig. 3.** Effects of sodium arsenite on gene transcription and readthrough transcripts. **a** Volcano plot comparing nascent transcription of genes of arsenite and mock-treated U2OS cells. Red color indicates induction, blue indicates repression. **b** Gene set enrichment analysis (GSEA) showing two top upregulated and two top downregulated gene sets. **c** Similar to heat shock treatment, arsenite induce transcription of the *HSPH1* gene. Reads from mock-treated cells in blue and heat shock treated cells in yellow. **d** Examples of four genes showing arsenite-induced readthrough transcripts (yellow).

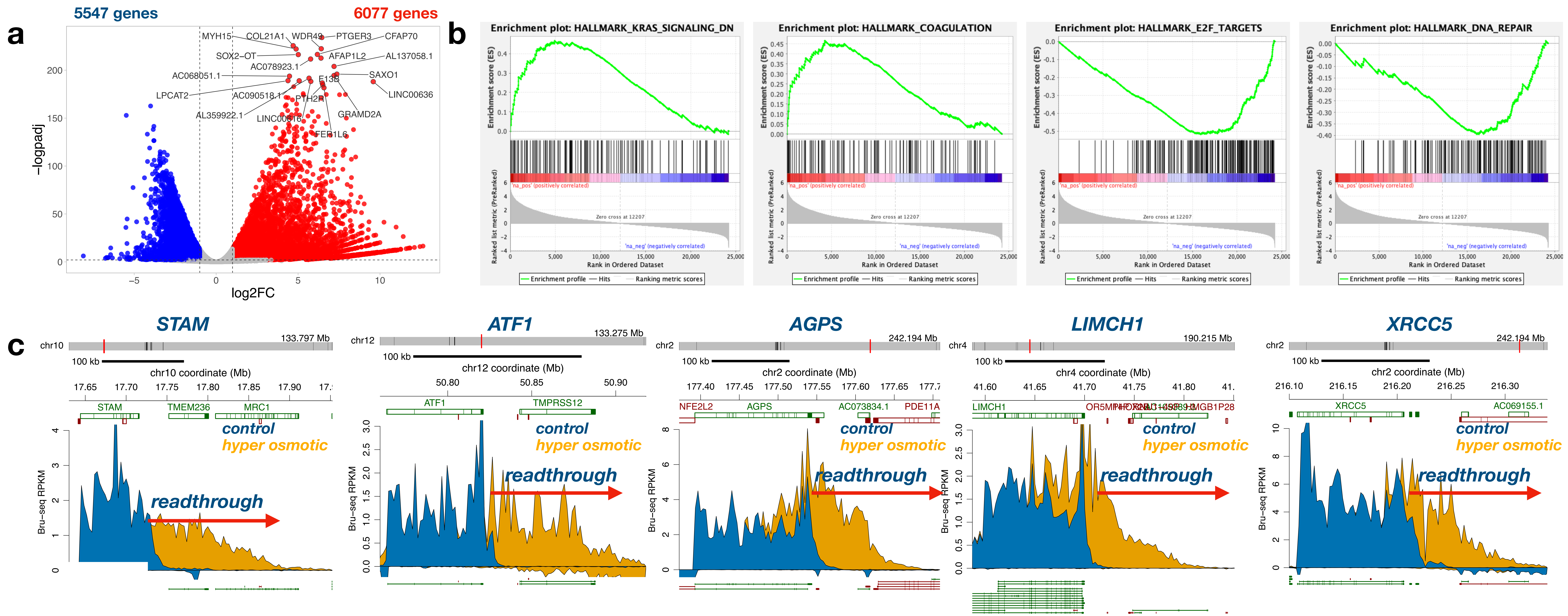


**Supplementary Fig. 4.** Effects of hyperosmotic stress on gene transcription and readthrough transcripts. **a** Volcano plot comparing nascent transcription of genes from cells exposed to hyper osmotic stress and from mock-treated U2OS cells. Red color indicates induction, blue indicates repression. **b** Gene set enrichment analysis (GSEA) showing two top upregulated and two top downregulated gene sets. **c** Examples of five genes showing hyper osmotic stress-induced readthrough transcripts (yellow).

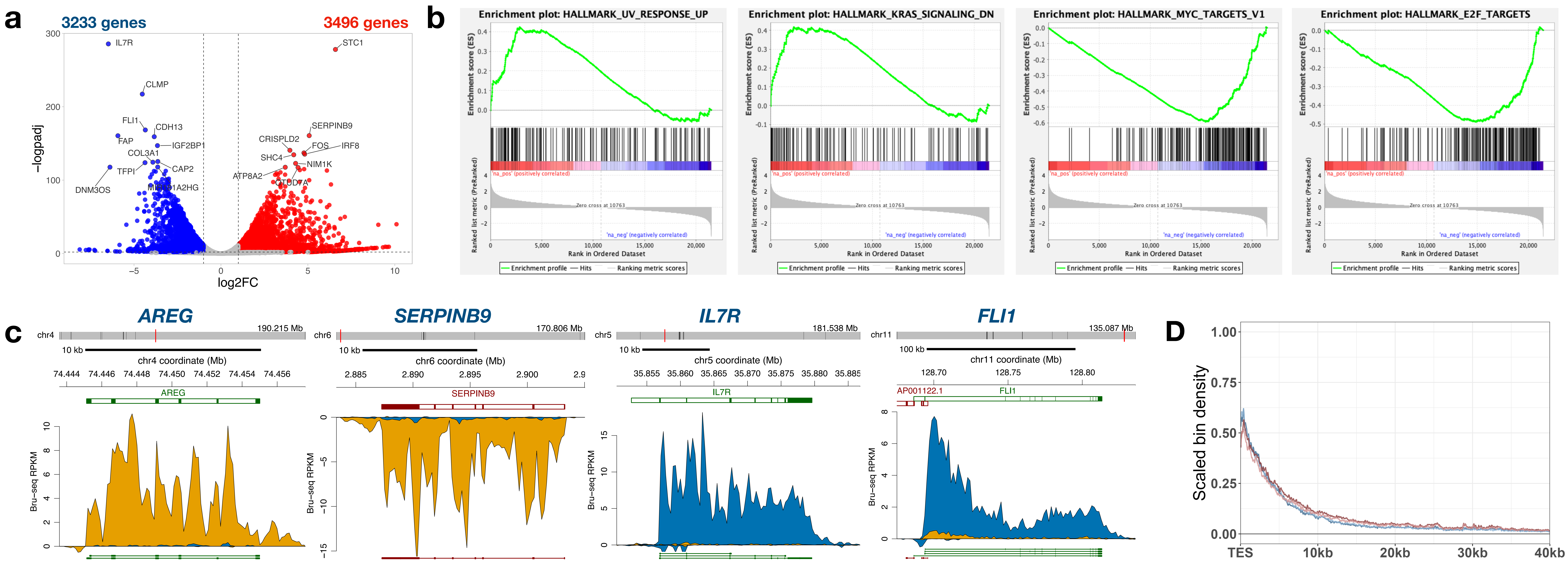


**Supplementary Fig. 5.** Effects of the HDAC inhibitor vorinostat on gene transcription and readthrough transcripts. **a** Volcano plot comparing nascent transcription of genes from HOS cells exposed to vorinostat and from mock-treated HOS cells. Red color indicates induction, blue indicates repression. **b** Gene set enrichment analysis (GSEA) showing two top upregulated and two top downregulated gene sets. **c** Examples of genes showing strong induction (*AREG* and *SERPINB9*) or repression (*IL7R* and *FLI1*) following vorinostat treatment. **d** No significant change in the scaled amount of readthrough transcripts was observed following vorinostat treatment.

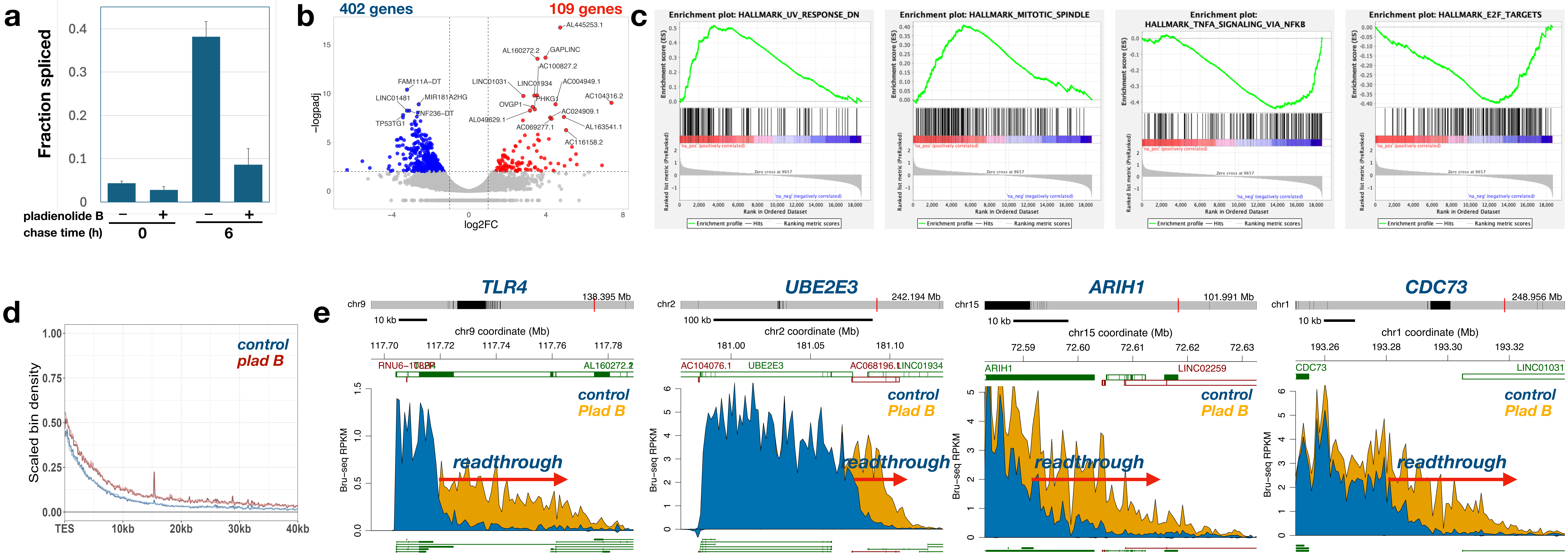


**Supplementary Fig. 6.** Effects of the splicing inhibitor plad B on gene transcription and readthrough transcripts. **a** Validation of splicing inhibition by plad B on HF1 fibroblasts showing suppression of the fraction of reads spliced both at 0 and 6h of chase. **b** Volcano plot comparing nascent transcription of genes from HF1 cells exposed to plad B and from mock-treated HF1 cells. Red color indicates induction, blue indicates repression. **c** Gene set enrichment analysis (GSEA) showing two top upregulated and two top downregulated gene sets. **d** Inhibition of splicing by plad B showed an increase in the scaled amount of readthrough transcripts compared to mock-treated control cells. **e** Examples of genes showing increases in readthrough transcripts following treatment with plad B.

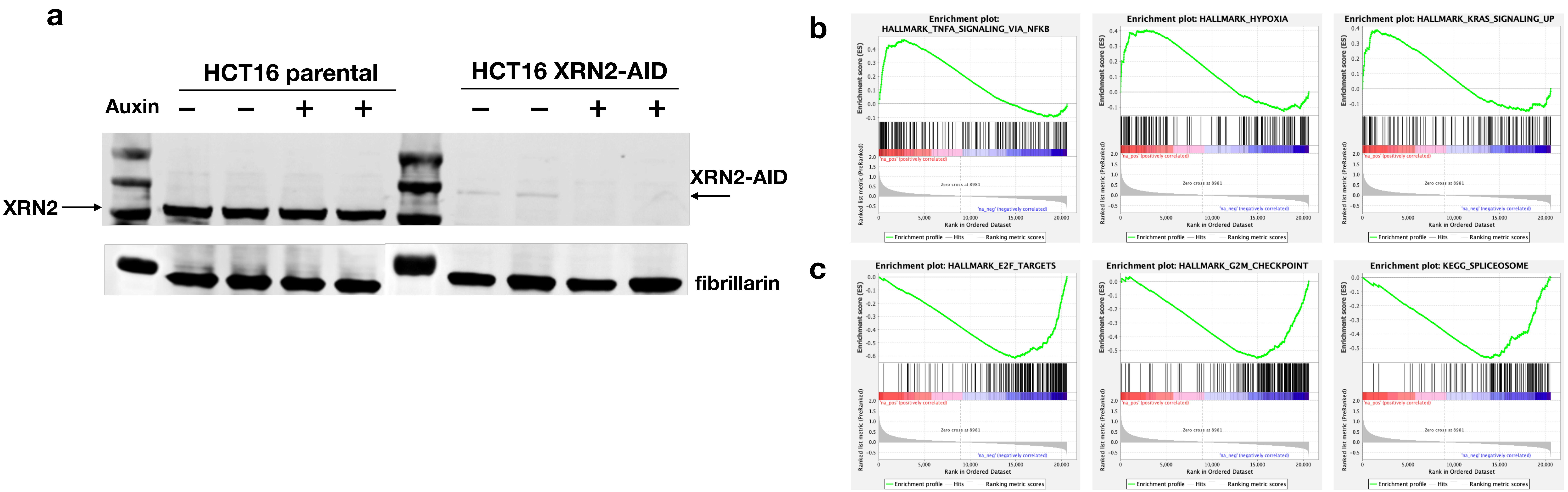


**Supplementary Fig. 7.** Auxin-mediated degradation of XRN2. **a** Western blot of parental (wt) and XRN2 degree cells either control treated or treated with auxin for different amounts of time. Fibrillarin was used as a loading control. **b** GSEA gene sets up regulated after x hours of auxin treatment. **c** GSEA gene sets down regulated by depletion of XRN2 by auxin.

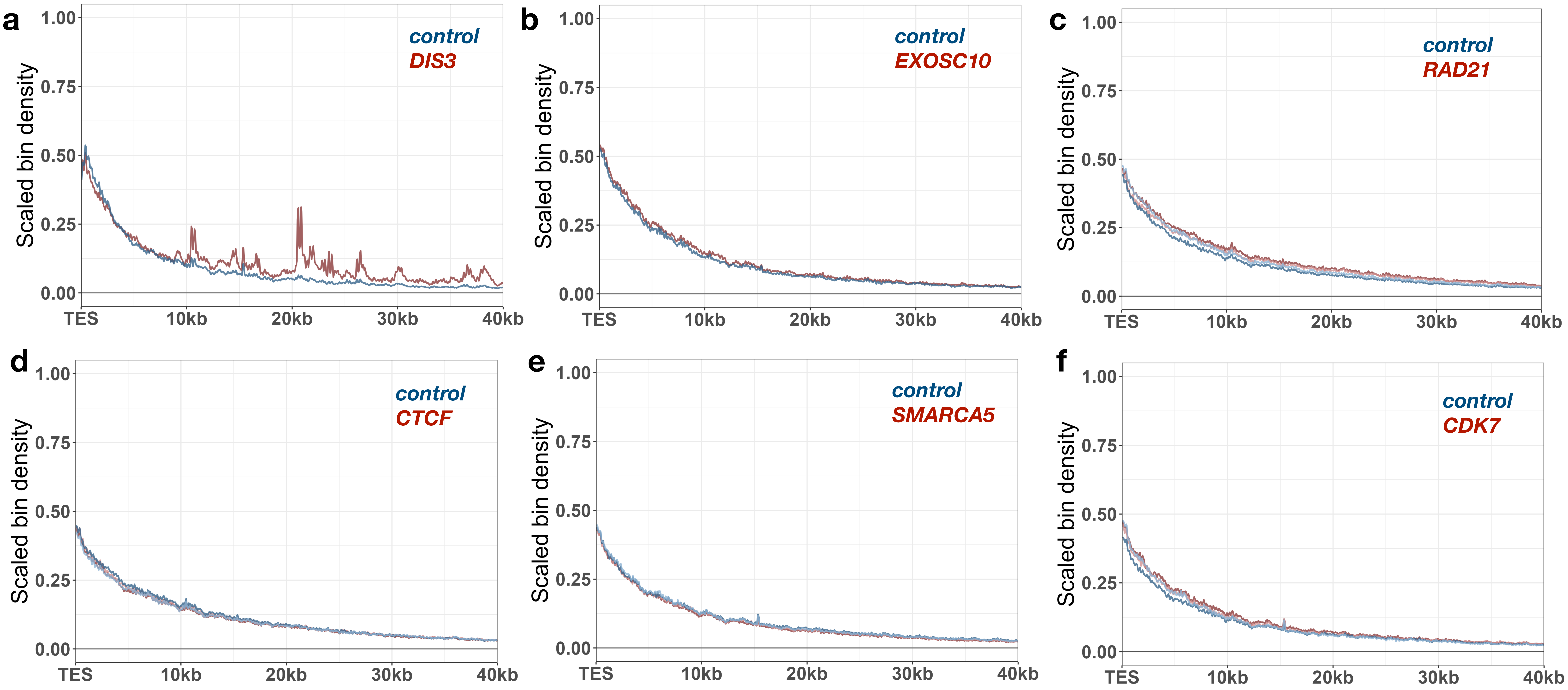


**Supplementary Fig. 8.** No effects of auxin-mediated depletion of various proteins on readthrough transcripts. Depletion of RNA exosome subunits DIS3 **a** and EXOSC10 **b**, chromatin organizers RAD21 **c** and CTCF **d,** chromatin remodeler SMARCA5 **e** or the kinase CDK7 **f**.

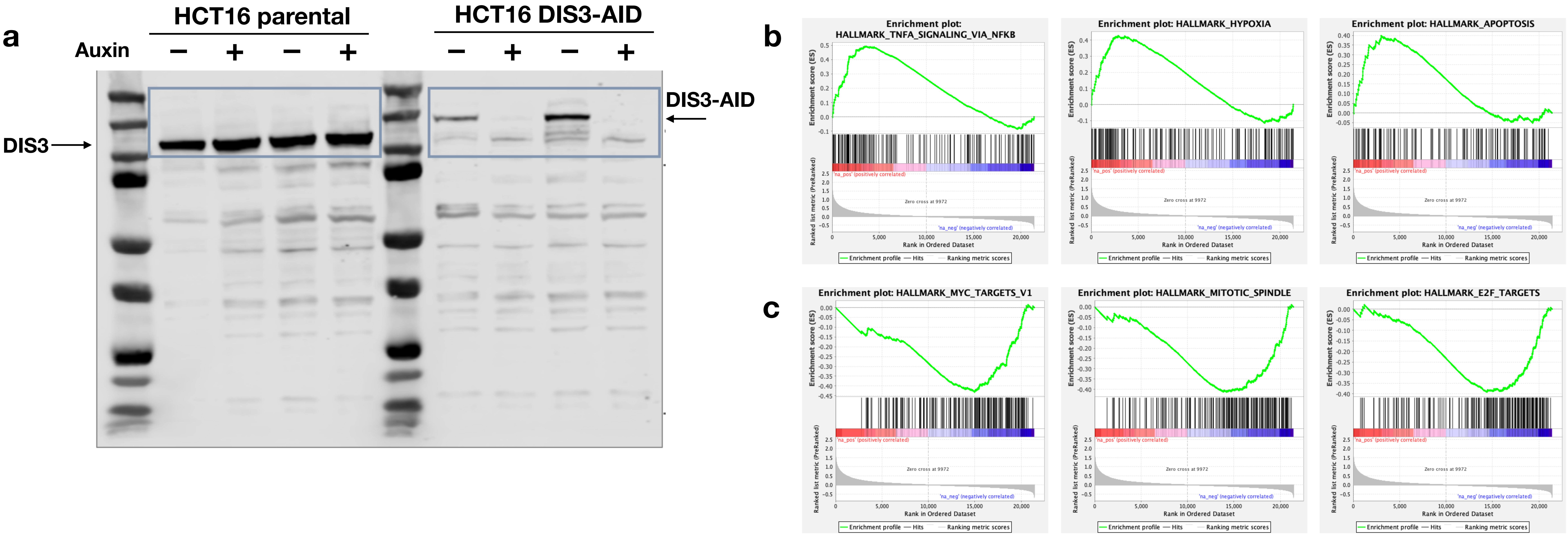


**Supplementary Fig. 9.** Auxin-mediated degradation of DIS3. **a** Western blot of parental (wt) and DIS3 degron cells either control treated or treated with auxin for 2 hours. **b** GSEA gene sets up regulated after 2 hours of auxin treatment. **c** GSEA gene sets down regulated by auxin treatment.

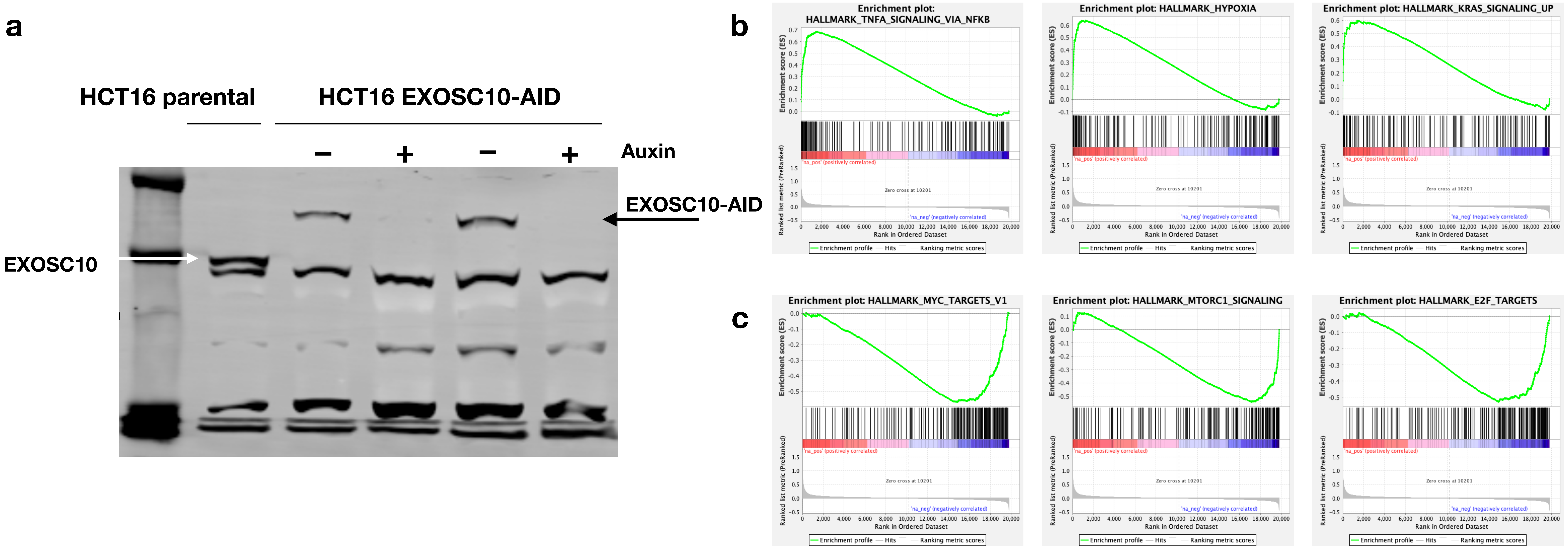


**Supplementary Fig. 10.** Auxin-mediated degradation of EXOSC10. **a** Western blot of parental (wt) and EXOSC10 degron cells either control treated or treated with auxin for 2 hours. **b** GSEA gene sets up regulated after 2 hours of auxin treatment. **c** GSEA gene sets down regulated by auxin treatment.

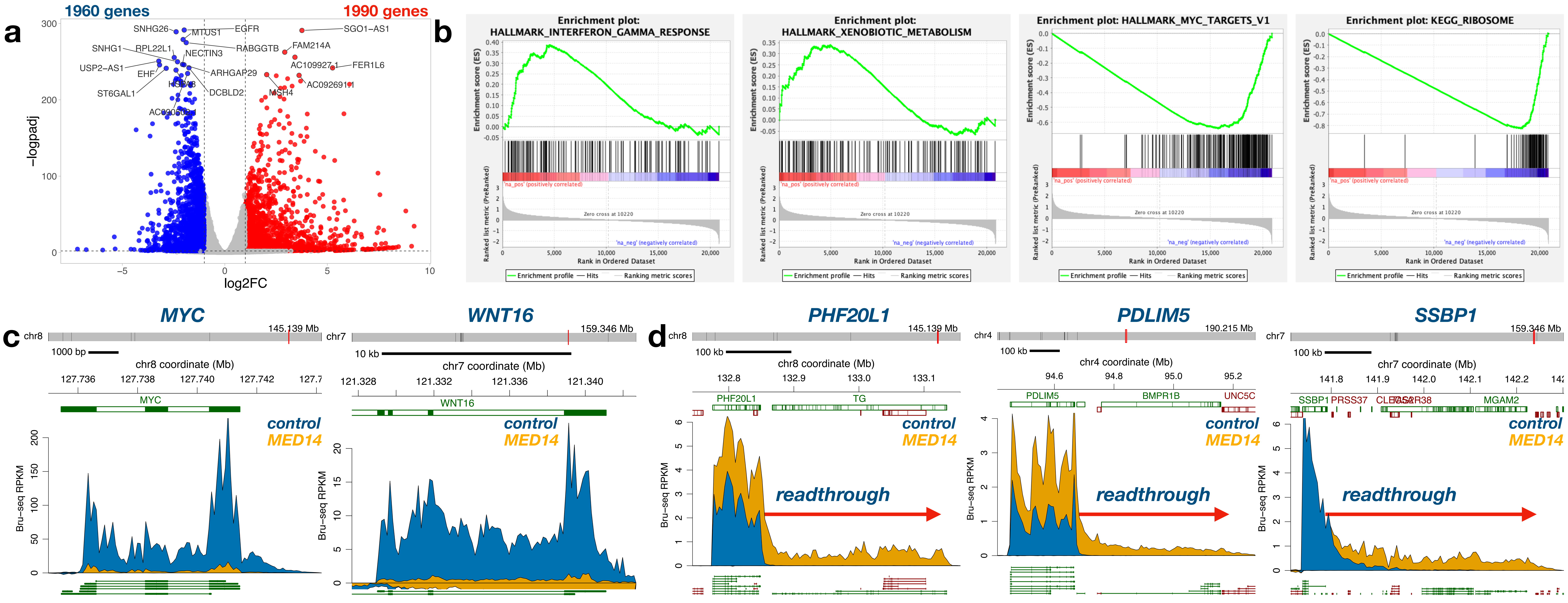


**Supplementary Fig. 11.** Effects of depletion of MED14 on gene transcription and readthrough transcripts. **a** Volcano plot comparing nascent transcription of genes of MED14-depleted or control HCT116 cells. Red color indicates induction, blue indicates repression. **b** Gene set enrichment analysis (GSEA) showing two top upregulated and two top downregulated gene sets. **c** Depletion of MED14 leads to repression of *MYC* and *WNT16* transcription. Reads from mock-treated cells in blue and MED14-depleted cells in yellow. **d** Examples of three genes showing MED14-depletion mediated readthrough transcripts (yellow).

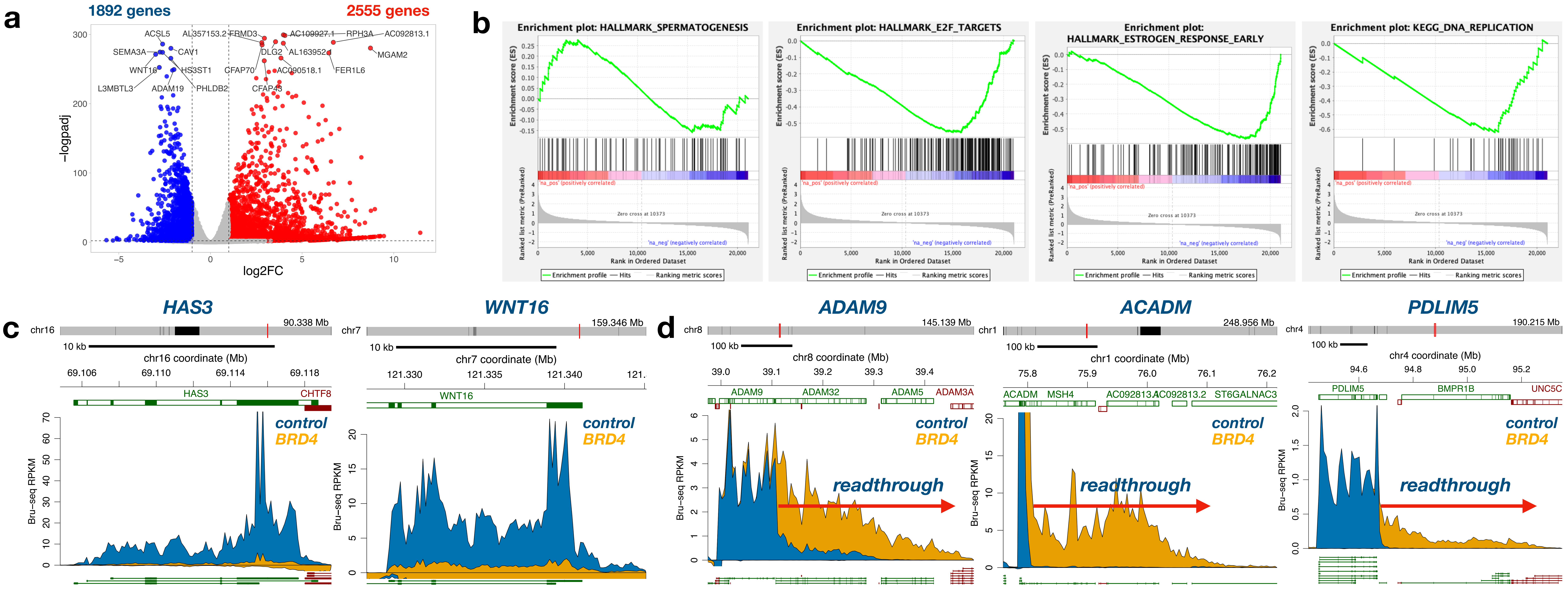


**Supplementary Fig. 12.** Effects of depletion of BRD4 on gene transcription and readthrough transcripts. **a** Volcano plot comparing nascent transcription of genes of BRD4-depleted or control HCT116 cells. Red color indicates induction, blue indicates repression. **b** Gene set enrichment analysis (GSEA) showing two top upregulated and two top downregulated gene sets. **c** Depletion of BRD4 leads to repression of *HAS3* and *WNT16* transcription. Reads from mock-treated cells in blue and MED14-depleted cells in yellow. **d** Examples of three genes showing BRD4-depletion mediated readthrough transcripts (yellow).

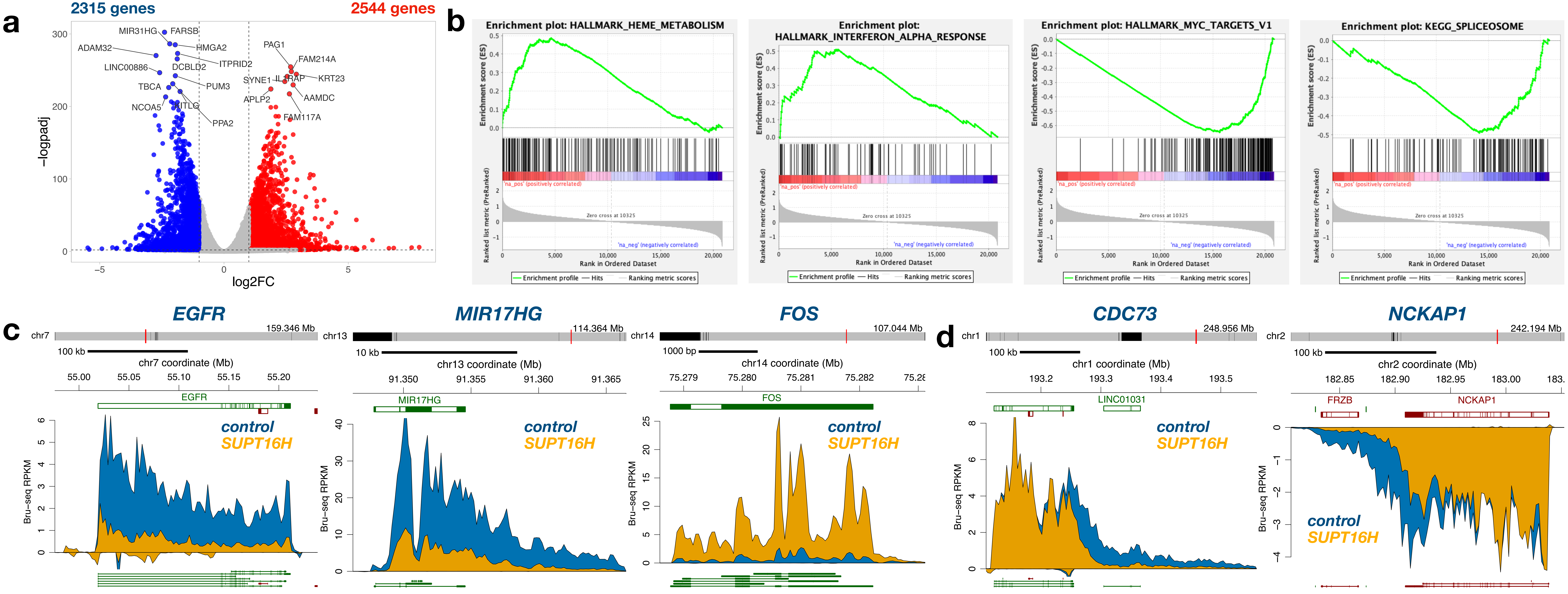


**Supplementary Fig. 13.** Effects of depletion of SUPT16H on gene transcription and readthrough transcripts. **a** Volcano plot comparing nascent transcription of genes of SUPTH16H-depleted or control HCT116 cells. Red color indicates induction, blue indicates repression. **b** Gene set enrichment analysis (GSEA) showing two top upregulated and two top downregulated gene sets. **c** Depletion of SUPT16H leads to repression of *EGFR* and *MIR17HG* and induction of *FOS* transcription. Reads from mock-treated cells in blue and SUPT16H-depleted cells in yellow. **d** Examples of two genes showing SUPT16H-depletion mediated suppression of readthrough transcripts (yellow).

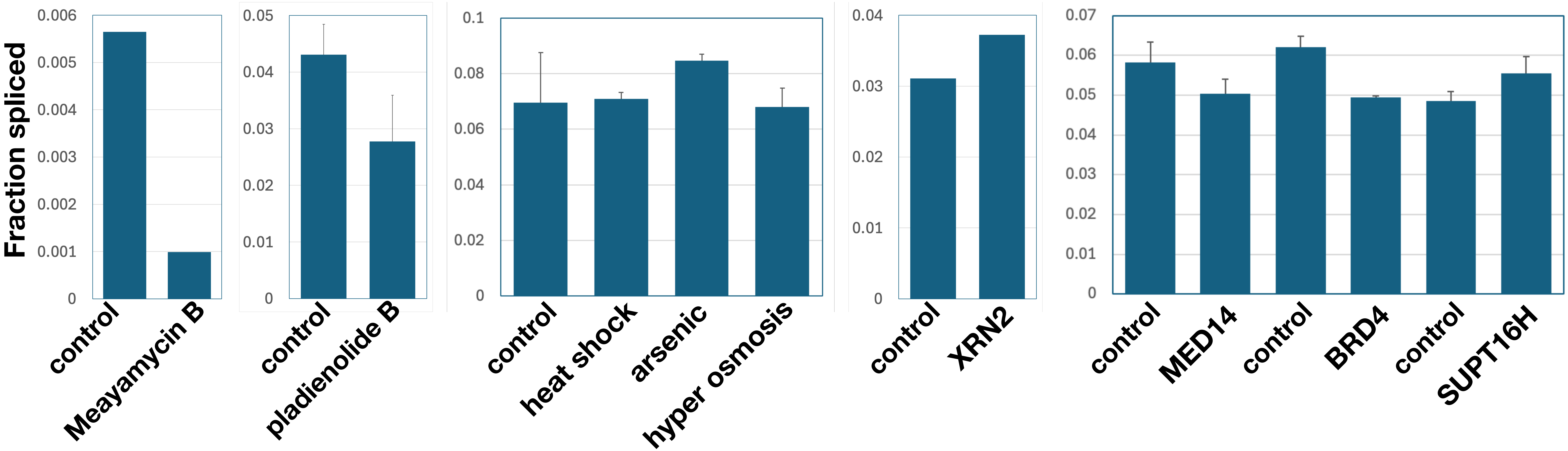


**Supplementary Fig. 14.** Splicing is inhibited by Meayamycin and plad B but not by other stressors or protein depletions. The fraction spliced was calculated and compared between control and treated samples.
